# Direct RNA sequencing of the *Escherichia coli* epitranscriptome uncovers alterations under heat stress

**DOI:** 10.1101/2024.07.08.602490

**Authors:** Sebastian Riquelme Barrios, Leonardo Vasquez Camus, Siobhan A. Cusack, Korinna Burdack, Dimitar Plamenov Petrov, G. Nur Yeşiltaç-Tosun, Stefanie Kaiser, Pascal Giehr, Kirsten Jung

## Abstract

Modifications of RNA, known as the epitranscriptome, affect gene expression, translation, and splicing in eukaryotes, with implications for developmental processes, cancer, and viral infections. In prokaryotes, regulation at the level of the epitranscriptome is still poorly understood. Here, we used Nanopore direct RNA sequencing of *Escherichia coli* to study RNA modifications and their changes under heat stress. With a single sequencing reaction, we detected most known modification types in rRNA, tRNA, and mRNA. RNA sequencing was complemented by a multifaceted approach that included mass spectrometry, deletion mutants, single-nucleotide PCR, and *in vitro* methylation. Known m^5^C and m^6^A sites in the rRNA were confirmed, but these types of modifications could not be localized in the mRNA. In response to heat stress, levels of m^5^C, m^6^A, and m^6,6^A increased in the 16S rRNA. Sequencing and mass spectrometry data demonstrated a decrease in tRNA modification abundance in the anticodon loop at 45 °C. In general, mRNA modifications at 37 °C were enriched in the coding regions of genes associated with general metabolism and RNA processing, which shifted to genes involved in cell wall synthesis and membrane transport under heat stress. This study provides new insights into the complexity of post-transcriptional regulation in bacteria.

**GRAPHICAL ABSTRACT:** 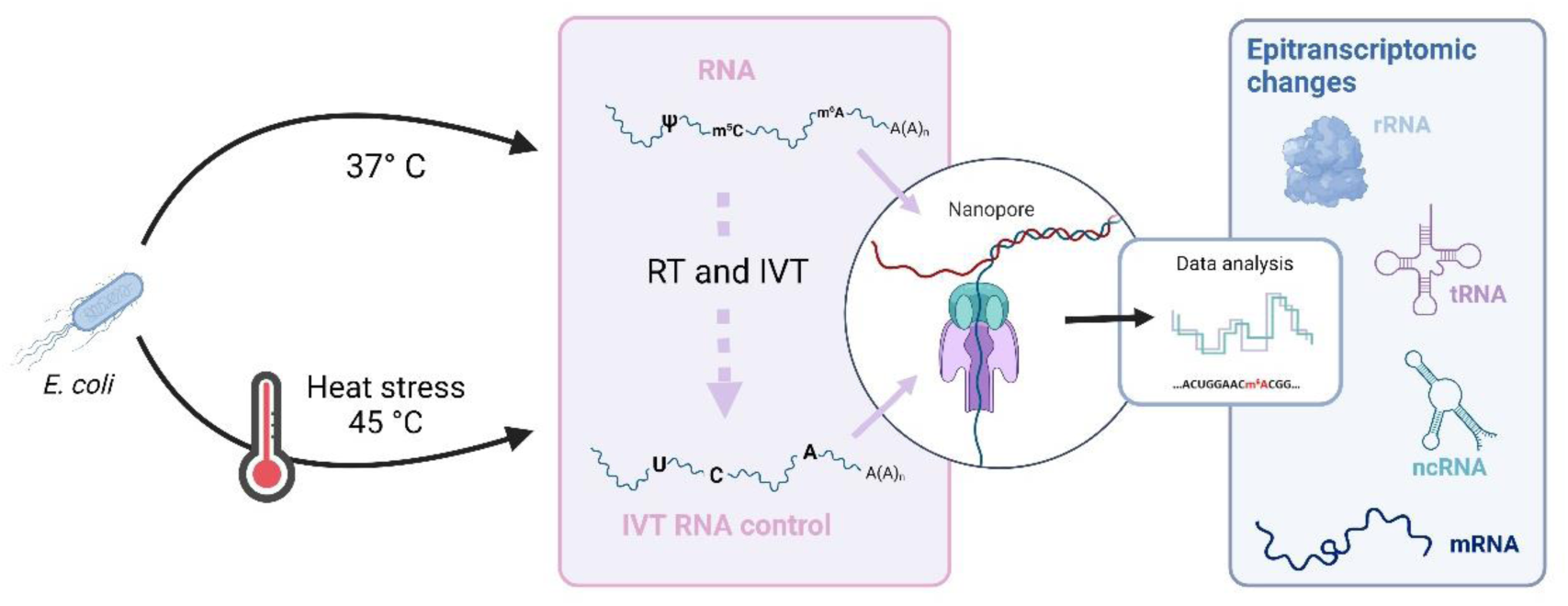

## INTRODUCTION

RNA molecules are subject to chemical modifications similar to those found in proteins and DNA. In prokaryotes, modifications in the ribosomal RNA (rRNA) and transfer RNA (tRNA) have already been characterized in detail. For example, rRNA modifications are known to play a key role in the catalytic activity and assembly of ribosomes, and the absence of such modifications can alter translation accuracy (1–4). tRNAs are also highly modified, particularly in the D- and TΨ-loops and in the anticodon region. These modifications affect the tRNA structure (5), mediate interactions with aminoacyl tRNA synthetases and elongation factor thermo unstable (EF-Tu) (6,7), and influence the affinity of the anticodon region for specific codons (8). Previous results from our laboratory have demonstrated the presence of numerous modification types in *E. coli* mRNA, which vary between growth stages and in response to stress conditions (9). Despite these earlier findings, bacterial mRNA modifications remain to be explored in detail; their roles, abundance, and distribution are still largely unknown.

Over the past decade, several platforms have been developed with the capacity to directly sequence native RNA molecules, providing new approaches to elucidate the epitranscriptome. Direct RNA sequencing (DRS) using Oxford Nanopore Technologies (ONT) platforms not only provides information about nucleotide (nt) sequences, but also about nt modifications. Modified bases produce slightly different electrical signals than canonical (i.e., unmodified) bases when the RNA strand translocates through the nanopore. Multiple experimental strategies and bioinformatics tools have been developed to identify RNA modifications from ONT’s DRS data. Some tools compare DRS signals between pairs of samples (e.g., wild-type [WT] vs. gene knockout [KO] or WT vs. *in vitro* transcribed [IVT] RNA) to identify signals corresponding to modifications (10–13), whereas other approaches are based on the evaluation of individual sample types (14,15). Although protocols have been developed to evaluate DRS data from bacterial samples (16) and the signals produced by bacterial rRNA modifications have been analyzed (17), comprehensive studies of regulation at the level of the epitranscriptome are still lacking.

In *E. coli*, detailed studies have been conducted for the detection of pseudouridine (Ψ) and *N*^6^-methyladenosine (m^6^A) (18,19). A protocol called Pseudo-Seq that combines carbodiimide N-cyclohexyl-N′-(2-morpholinoethyl)carbodiimide methyl-p-toluene sulfonate (CMC) treatment, size selection for truncated reverse-transcribed cDNA, and high-throughput sequencing revealed that RluA, previously characterized as a tRNA- and 23S rRNA-specific Ψ-synthase, is responsible for modifications of at least 31 different mRNA positions (18). Other rRNA Ψ-synthases, such as RluC and RluD, also modify some mRNA positions (18). Notably, *fimA* transcripts (which encode the major subunit of the pilus) are less abundant in a Δ*rluA* mutant (18).

m^6^A is the most prevalent mRNA modification in eukaryotes and has roles in translation, degradation, splicing, nuclear export, and mRNA turnover (20). In Gram-negative bacteria, such as *E. coli* and *Pseudomonas aeruginosa*, the ratio of m^6^A to unmodified A bases has been calculated as ∼0.2%, similar to the ratios found in eukaryotic cells (19). Immunoprecipitation of m^6^A followed by Illumina sequencing has revealed m^6^A modifications in the mRNA of bacterial genes related to metabolism, transport, gene regulation, the cell envelope, ribosomes, stress responses, and nucleic acid synthesis (19). m^6^A consensus motifs have been proposed in *E. coli* and *P. aeruginosa* (UGCCAG and GGCCAG, respectively). Furthermore, a recent study using a combination of DRS and antibodies for m^6^A RNA immunoprecipitation (MeRIP-seq) describes 75 high-confidence m^6^A sites in the mRNA corresponding to 21 *E. coli* genes (21). However, no m^6^A methyltransferase for mRNA is currently known (19,21).

In nature, microbes are frequently exposed to various environmental stresses, including sudden changes in growth temperature (e.g., heat stress). In *E. coli*, heat stress triggers a well-characterized response involving the induction of heat-stress proteins to counteract protein denaturation and prevent protein aggregate accumulation (22,23). Mechanisms for DNA repair (24) and reactive oxygen species mitigation (25) are also activated at elevated temperatures, as are modifications that help stabilize the tRNA to facilitate temperature adaptation. Specifically, levels of modifications such as 4-thiouridine (s^4^U), 2’-O-methylguanosine (Gm) at position 18, and 7-methylguanosine (m^7^G) at position 46 increase at high temperatures (26), whereas levels of modifications at position 32 decrease (26,27). In the tRNA T-loop, the presence of 5-methyluridine (m^5^U) at position 54 (catalyzed by TrmA) and Ψ55 (produced by TruB) enhances stacking interactions (28,29); the absence of these modifications and of Gm18 (introduced by TrmH) impairs growth under heat stress (4,30).

Heat stress also impacts modifications in the rRNA. For example, 5-hydroxycytidine (ho^5^C) at position 2501 of the 23S rRNA in *E. coli* diminishes after one hour of exposure to 45 °C (31). *In vitro* studies suggest that a high ho^5^C modification level almost inhibits protein biosynthesis (31). Similarly, mutants for *rlmE*, which encodes the heat shock protein responsible for methylation of 2’-O-ribose U2552 in the 23S rRNA, show alterations in the ribosome profile and a severe growth defect (32). Mutants of the methyltransferases that are responsible for 5-methylcytidine (m^5^C) at position 1407 and *N*^6^,*N*^6^-dimethyladenosine (m^6,6^A) at positions 1518 and 1519 have also previously demonstrated decreased responses to heat stress (33). Although changes in specific types of tRNA and rRNA modifications at individual positions have been studied, a comprehensive analysis of all modification types simultaneously under stress conditions has not yet been conducted, and stress-induced changes in modification levels have not been assessed in *E. coli* mRNA.

To address knowledge gaps regarding prokaryotic RNA modifications and their changes in response to stress, we here established a systematic DRS approach to thoroughly and simultaneously study numerous modification types in the rRNA, tRNA, and mRNA of *E. coli* K-12 (WT), an IVT RNA control, and an m^6^A methyltransferase KO mutant. This approach was complemented by mass spectrometry, single nucleotide qPCR, and *in vitro* methylation to localize m^5^C and m^6^A and other modifications in the mRNA. These approaches yielded a uniquely robust data set, enabling the first comprehensive analysis of the epitranscriptomic landscape in *E. coli* and its changes under heat stress. Our study not only advances understanding of the importance of RNA modifications in this model bacterium but establishes a blueprint for future epitranscriptomic analyses in prokaryotes.

## MATERIALS AND METHODS

### Strains and growth conditions

Experiments were conducted with the WT *E. coli* strain MG1655. Bacteria were grown in lysogeny broth (LB) at 37 °C with 200 rpm shaking to an optical density at 600 nm (OD_600_) of 0.5. Untreated cells were retained at 37 °C until they reached an optical density at 600 nm (OD_600_) of ∼0.8–1; for heat-shock treatment, 3.52 ml of 90 °C LB was added to 20 ml of culture, which was then incubated at 45 °C. Cells from both groups were collected 30 min later. The single mutants Δ*rlmF* and Δ*rlmJ* and the double mutant Δ*rlmF*Δ*rlmJ* were generated with in-frame deletion using an approach similar to a previously described method (34). Briefly, ∼500-nt regions upstream and downstream of the gene to be deleted were amplified in two overlapping PCRs; purified; ligated into the suicide vector pNPTS138-R6KT using Gibson assembly; and transferred into *E. coli* DH5α λpir. The vector was then transferred in a second transformation into the diaminopimelic acid (DAP) auxotrophic *E. coli* WM3064 donor strain. The non-replicative plasmid was transformed into WT MG1655 via conjugative mating on LB agar with the addition of 300 μM DAP to site-specifically integrate the plasmid into the genome. Bacteria with a chromosomally integrated plasmid were selected on LB agar with kanamycin sulfate. Strains with the plasmid integrated were selected the following day by growth on LB agar with 10% sucrose (w/v) at 30 °C. Sucrose-resistant kanamycin-sensitive strains were then examined for deletion of the desired gene using single-clone PCR and sequencing. Δ*trmM* mutants were generated using the Quick & Easy *E. coli* Gene Deletion Kit (Gene Bridges GmbH, Heidelberg, Germany), which uses the RED®/ET® recombinase system (35). Primers used for mutant construction are listed in **Supplementary Table 1**.

### Total RNA isolation

RNA was isolated using the PCI protocol (36) with some modifications (9). Specifically, bacterial cultures were mixed with phenol and ethanol to final concentrations of 1% (v/v) and 20% (v/v), respectively, then flash-frozen in liquid nitrogen. Thawed samples were centrifuged at 16,000 × g for 10 min and the resulting pellets were resuspended in 500 µL of ice-cold sodium acetate buffer (20 mM, pH 5.2) with 1 mM EDTA (AE buffer). Extracted RNA was treated with RNase-free DNase I (New England Biolabs [NEB], Ipswich, MA, USA) following the manufacturer’s protocol. RNA quality and integrity were assessed via chip gel electrophoresis with a 2100 Bioanalyzer and an RNA Nano chip kit (Agilent Technologies, Santa Clara, CA, USA).

### tRNA/rRNA removal and mRNA enrichment

To remove tRNAs, total RNA samples were purified using the RNA Clean & Concentrator Kit (Zymo Research, Irvine, CA, USA) following the manufacturer’s protocol for exclusion of RNA below 200 nt. RNA quality was confirmed by electrophoresis using the 2100 Bioanalyzer (Agilent Technologies, Santa Clara, CA, USA). Oligo-based rRNA depletion was then performed with the pan-Bacteria riboPOOL rRNA Depletion Kit (siTOOLS, Planegg, Germany) according to the manufacturer’s instructions. Depletion efficiency was assessed using Bioanalyzer mRNA chip electrophoresis (with Pico and Nano chips) and reverse transcription (RT) quantitative PCR (qPCR) as described below to amplify 23S and 16S rRNA and the mRNA marker *recA*. Ribosome (16S and 23S) transcript abundance was measured both before and after the depletion protocol was applied and normalized to *recA* abundance in each sample. Ribosome depletion was then calculated as the percentage of normalized 23S and 16S rRNAs remaining after the depletion protocol. Primers for RT-qPCR are shown in **Supplementary Table 2**.

### RT-qPCR

cDNA was reverse transcribed from purified (DNase-treated) RNA samples. RT was carried out with the iScript™ Advanced cDNA Synthesis Kit (Bio-Rad, Hercules, CA, USA) on 500 ng to 1 µg RNA per sample according to the manufacturer’s protocol. The resulting RT product was diluted four times and 4 μl per sample was mixed with 1 μl of primer mix (containing 10 μM of each primer) and 5 μl of SsoAdvanced™ Universal SYBR® Green Supermix (Bio-Rad, Hercules, CA, USA). Amplification was conducted on the CFX96^TM^ Real-Time System (Bio-Rad, Hercules, CA, USA) using 16S rRNA as the internal control; a No-RT negative control (RNA sample without RT enzyme) and a No-template negative control (water and qPCR mix with no template) were both included in every experiment. Relative gene expression was calculated as the abundance of the gene of interest normalized to that of the 16S rRNA using the X^- ΔCt^ method, where X corresponds to primer-specific amplification efficiency. RT-qPCR primer sequences are shown in **Supplementary Table 2**.

### Mass spectrometry (MS) analysis of RNA methylation

Absolute quantitative analyses using isotope dilution MS were performed as previously described (9). Briefly, RNAs (100–300 ng per sample) were enzymatically hydrolyzed to nucleosides by incubation with a mixture of benzonase (2 U), snake venom phosphodiesterase (0.2 U), and calf intestine phosphatase (2 U) for 2 h at 37 °C in a buffer containing Tris-HCl (5 mM, pH 8), deoxycoformycine (1 µg), and tetrahydrouridine (5 µg). For calibration, the nucleosides cytidine (C), uridine (U), guanosine (G), adenosine (A) (Sigma-Aldrich, St. Louis, MO, USA), and m^6^A (Carbosynth, Staad, Switzerland) were each mixed to a final concentration of 100 nM (canonical nucleosides) or 5 nM (m^6^A). This stock calibration solution was serially diluted 1:2 to a total of 12 calibration levels. Prior to analysis, the improved gen ^13^C/^15^N stable isotope-labeled internal standard (SILIS) from *Saccharomyces cerevisiae* tRNA (37) was added for absolute quantitative analysis. The resulting ribonucleotide mixture was separated using a Synergy Fusion RP column (Phenomenex, Torrance, CA, USA) with a 2.5-μm particle size, 100-Å pore size, 100-mm length, and 2-mm inner diameter on an Agilent 1290 Infinity II series ultra-high-performance liquid chromatography (UHPLC) instrument. Mobile phase A was 5 mM ammonium acetate adjusted to pH 5.3 with glacial acetic acid and mobile phase B was pure acetonitrile. Gradient elution began with 100% A for 1 min, increased to 10% B at 4 min and 40% B at 7–8 min, then returned to the starting condition of 100% A for an additional 2.5 min. The flow rate was 0.35 mL/min and the column temperature was 35 °C. For MS measurements, an Agilent 6470 Triple Quadrupole MS instrument set to dynamic multiple reaction monitoring mode was used. MS was conducted in positive ion mode with the following parameters: skimmer voltage of 15 V, cell accelerator voltage of 5 V, N_2_ gas temperature of 230 °C, N_2_ gas flow of 6 L/min, sheath gas (N_2_) temperature of 400 °C with a flow of 12 L/min, capillary voltage of 2500 V, nozzle voltage of 0 V, and nebulizer at 40 psi. A list of monitored mass transitions is given in **Supplementary Table 3**. Data analysis was performed using MassHunter Quantitative software to yield molar amounts of analytes. Absolute m^6^A abundance was calculated as the abundance of modified As per 1000 total As.

### Purification of RlmJ and RlmF

*rlmJ* (b3499) and *rlmF* (b0807) were cloned into the pET28 vector (bearing a C-terminal His6-tag). *rlmJ* was amplified from the *E. coli* genome and ligated into the pET28 vector using Gibson assembly. *rlmF* was amplified via PCR, then the amplicon was cut with NcoI and HindIII (NEB, Ipswich, MA, USA) and ligated into the vector pET28, which had also been digested with NcoI and HindIII. The sequences were verified with sequencing. Primers used for cloning are shown in **Supplementary Table 4**. *E. coli* BL21 (DE3) pLysS competent cells (Novagen, Darmstadt, Germany) containing the pET28-based expression system were cultured in LB at 37 °C to an OD_600_ of 0.5, then gene expression was induced by addition of 0.5 mM isopropyl β-D-thiogalactoside (IPTG) for 3 h at 37 °C to a final OD_600_ of ∼2. Cell pellets were lysed by sonication in buffer containing 20 mM Tris-HCl (pH 7.0), 200 mM NaCl, 5% glycerol (v/v), and 5 mM β-mercaptoethanol (BME) as previously reported (38). The lysate was centrifuged for 10 min at 5,000 x g, then for 30 min at 70,000 x g. The resulting supernatant was incubated with Ni-NTA Agarose (QIAGEN, Hilden, Germany) for 45 min at 4 °C, then washed with 30 volumes of 20 mM Tris-HCl (pH 7.0), 500 mM NaCl, 5% glycerol (v/v), 5 mM BME, and 10 mM imidazole. Enzymes were eluted with 20 mM Tris-HCl (pH 7.0), 500 mM NaCl, 5% glycerol (v/v), 5 mM BME, and 100 mM EDTA. Proteins were concentrated using Amicon® (Merck, Darmstadt, Germany) and stored at 4 °C for up to one month.

### *In vitro* transcription for methyltransferase activity assays

Following the manufacturer’s protocols, RNA was synthesized from a DNA template (**Supplementary Table 5**) using the HiScribe T7 High Yield RNA Synthesis Kit (NEB, Ipswich, MA, USA) and treated with DNase I (NEB, Ipswich, MA, USA) to eliminate DNA. The resulting RNA was purified and concentrated with a Monarch RNA Clean-up Kit (NEB, Ipswich, MA, USA), quantified, and stored at -80 °C prior to further use. For methylation reactions, IVT RNA (50–200 pmol) was heated for 1 min at 95 °C, then cooled on ice. The reaction was carried out for 10 min at 37 °C in a buffer containing 20 mM potassium phosphate (pH 6.5), 50 mM NaCl, 5 mM BME, 1 U/µl murine RNase inhibitor (NEB, Ipswich, MA, USA), and 6 μg of pure enzyme (RlmJ/RlmF) or bovine serum albumin (NEB, Ipswich, MA, USA) as a control (38). Finally, 0.5 µM S-adenosyl-L-[methyl-^3^H]-methionine (SAM) (5–15 Ci/mmol; American Radiolabeled Chemicals, Inc., Saint Louis, MO, USA) was added to a total volume of 50 μL to start the reaction. After 10 min, the reaction was stopped using RNA binding buffer from the RNA Clean & Concentrator Kit (Zymo Research, Irvine, CA, USA) plus ethanol following the manufacturer’s protocol. Purified RNA was diluted in scintillation liquid, homogenized, and quantified in a Tri-Carb® 2910 TR scintillation counter (PerkinElmer, Waltham, MA, USA). Previously reported methylated sites in the 23S were used as positive controls. For RlmJ, the positive control was a 28-nt fragment containing the A2030 position of the 23S rRNA with an additional 5′-GG sequence (39). For RlmF, a 30-nt fragment from 1601 to 1631 of the *E. coli* 23S rRNA containing A1618 served as the positive control.

### SELECT method for m^6^A detection

The SELECT method was conducted as previously described (40,41). Briefly, total RNA from WT or IVT samples was hybridized with 40 nM up/down primers (**Supplementary Table 6**), then combined with 5 μM dNTPs and 2 μL of 10× rCutSmart buffer (NEB, Ipswich, MA, USA) in a total reaction volume of 17 μL. For positions in the 23S rRNA, 100 ng of RNA was used; reactions for all other positions included 500 ng of RNA. The mixture was annealed sequentially at 90, 80, 70, 60, and 50 °C for 1 min each, then at 40 °C for 6 min. Subsequently, 3 μL of a ligation mixture containing 0.01 U of Bst 2.0 DNA polymerase (NEB, Ipswich, MA, USA), 0.5 U of SplintR ligase (NEB, Ipswich, MA, USA), and 10 nmol ATP was added. The mixture was incubated at 40 °C for 20 min, then denatured at 80 °C for 20 min. RT-qPCR was conducted with 4 μL of the product using 200 nM of SELECT specific forward and reverse primers (**Supplementary Table 2**).

### IVT RNA control preparation

The IVT RNA control was prepared as previously described (42,43) with some modifications. Briefly, 50 ng of polyadenylated rRNA-depleted *E. coli* RNA was used as the starting material. VN and Strand-Switching primers from the cDNA-PCR Sequencing Kit (ONT #SQK-PCS109) were used with Maxima H Minus Reverse Transcriptase (Thermo Fisher Scientific, Waltham, MA, USA) for RT and strand switching with the following parameters: 90 min at 42 °C (one cycle), heat inactivation for 5 min at 85 °C (one cycle), then holding at 4 °C. The RT RNA sample (5 μL) was amplified using the LongAmp Taq 2× Master Mix (NEB, Ipswich, MA, USA) with IVT Nanopore T7 Fw and IVT Nanopore T7 Rv primers (**Supplementary Table 7**). The thermocycling conditions were as follows: initial denaturation for 30 s at 95 °C (one cycle), denaturation for 15 s at 95 °C (11 cycles), annealing for 15 s at 62 °C (11 cycles), extension for 5 min at 65 °C (11 cycles), and a final extension for 6 min at 65 °C (one cycle), followed by holding at 10 °C. The PCR product was treated with Exonuclease I (NEB, Ipswich, MA, USA) for 15 min at 37 °C, followed by 15 min at 80 °C to eliminate single-stranded products. DNA was purified using AMPure XP (Beckman Coulter, Brea, CA, USA) following the manufacturer’s instructions for the cDNA-PCR Sequencing Kit (ONT #SQK-PCS109). The PCR product was used as the template for IVT with the HiScribe T7 High Yield RNA Synthesis Kit (NEB, Ipswich, MA, USA). Samples were then treated with DNase I and cleaned with the Monarch RNA Cleanup Kit (NEB, Ipswich, MA, USA) following the manufacturer’s protocol. RNA was polyadenylated with *E. coli* Poly(A) Polymerase (NEB, Ipswich, MA, USA) and cleaned with the Monarch RNA Cleanup Kit (NEB, Ipswich, MA, USA). Ribosomal 23S and 16S samples were prepared by amplifying the appropriate sequences from genomic *E. coli* DNA and adding the T7 sequence to the forward primer (**Supplementary Table 7**). IVT was performed following the manufacturer’s protocol for the HiScribe T7 High Yield RNA Synthesis Kit (NEB, Ipswich, MA, USA). Polyadenylated RNA was added to the IVT RNA control at a ratio of 1:1000.

### Nanopore sequencing

mRNA-enriched samples and IVT RNA controls were analyzed via DRS using the MinION platform (ONT, Oxford, UK). The DNase-treated, tRNA-depleted RNA samples were polyadenylated with *E. coli* Poly(A) Polymerase (NEB, Ipswich, MA, USA) for 30 min following the manufacturer’s protocol. Polyadenylation levels were assessed with Bioanalyzer chip gel electrophoresis and through evaluation of a size shift increase of ∼500 bp for the 23S and 16S rRNAs. rRNA was depleted as described above. Nanopore libraries were prepared with ∼200 ng of polyadenylated *E. coli* mRNA or IVT RNA following the manufacturer’s protocol (ONT #SQK-RNA002). Each sample was sequenced on an individual MinION R9.4.1 flow cell.

### ONT data processing

Reads from ONT sequencing were basecalled using Guppy v6.5.7 (ONT, Oxford, UK) with the rna_r9.4.1_70bps_hac.cfg configuration file and the two biological replicates of each sample type combined to form a single sample each. The total numbers of sequenced reads and bases passing quality control during basecalling, average read length, and N50 length were calculated with SeqKit

(44). Average Q scores were calculated with NanoPlot (45). Minimap2 (v2.26) (46) was used for alignment to a modified version of the ensemble *E. coli* reference genome (Escherichia_coli_str_k_12_substr_mg1655_gca_000005845.ASM584v2) with duplicate tRNA and rRNA genes removed and to a transcriptome generated using the coordinates of transcriptional units, including 5′ and 3′ untranslated regions (UTRs) (47). Mapped reads were divided into the following RNA types as annotated in the EnsemblBacteria cDNA and ncRNA FASTA files for *E. coli* str. K-12 substr. MG1655 (GCA_000005845): tRNA; rRNA; protein coding; nontranslating coding sequence (CDS); pseudogene; and noncoding RNA (ncRNA), comprising small RNA (sRNA) and small interfering RNA (siRNA). Reads mapping to *ssrA* were classified as transfer-messenger RNA (tmRNA). The numbers of mapped reads and bases belonging to each RNA type and the numbers of unmapped reads and bases that passed quality control during basecalling were calculated with SeqKit (44). For further analyses, aligned reads were filtered using SAMtools (v1.17) (48) to remove secondary alignments, then quantified using Salmon (49) and analyzed with ELIGOS (11) to identify putative modifications of all types. Aligned reads were also indexed and preprocessed with Nanopolish (v0.14) (50) before analysis with CHEUI (51) for detection of only m^6^A and m^5^C and analysis with Nanocompore (52) and DRUMMER (13) (in exome mode) for detection of all modification types. With the exception of a minimum read count of 20 in Nanocompore, all programs were run with default parameters. For differentially expressed gene (DEG) analysis, read counts from Salmon were imported into R using the ‘tximport’ package (53), and DEGs were identified using DESeq2 (v1.38.3) (54). Note, only genes that were present in at least three samples at a depth of at least 10 reads were considered for analysis. Significant DEGs were obtained by filtering the results based on *p*-value (≤ 0.05), *q*-value (≤ 0.05), and log_2_(fold change) (≥ 2).

### Putative modification site analyses

#### Biotype annotations

Putative modification sites were classified as present within the tRNA, rRNA, ncRNA, CDS, 5′ UTR, 3′ UTR, or intergenic region using the annotated *E. coli* K-12 substr. MG1655 ASM584v2 reference genome (RefSeq #GCF_000005845.2) and the genomic coordinates of UTRs (47).

#### Thresholding

Precision–recall curves were generated for modifications in the rRNA and tRNA using a range of thresholds for each of several values generated by ELIGOS: the percent error of specific bases (ESB) in the WT sample (ESB test), ESB in the IVT control sample (ESB control), the odds ratio (OddR), and the adjusted *p*-value. Precision and recall were also calculated for all possible combinations of the threshold values tested for these four parameters. For these analyses, recall was calculated using known modification sites only. Precision was calculated considering known modification sites ± 10 nt within continuous regions of putative modification sites as true signals due to the known noise caused by modifications in Nanopore data. The final thresholds for the ESB test, ESB control, OddR, and *p*-value were selected by taking the combination of threshold values that resulted in the highest recall at a precision of ≥ 85% (i.e., error ≤ 15%). Where multiple values of a single parameter resulted in identical precision and recall values, the least restrictive threshold value was selected.

#### Modified region identification and characterization

The threshold values established as described above were applied to modification sites called by ELIGOS in all biotypes other than the tRNA and rRNA (namely the CDS, ncRNA, intergenic region, and 5′ and 3′ UTR). Due to the presence of known modification sites in the tRNA and rRNA, true modification sites could be distinguished from noise within regions of adjacent nt in which all sites were called as modified by ELIGOS; because this was not possible for the other gene biotypes, each group of adjacent nt in which every site was called as modified by ELIGOS was considered a single “signal region” in which one or more modifications were present. The distribution of modifications across a model gene transcript was determined by calculating the normalized relative location of each modification within the 5′ UTR, CDS, or 3′ UTR. The data were visualized with the R package ‘ggplot2’ (55).

### Statistical analyses

Differences between pairs of samples were assessed with an unpaired Student’s *t*-test. Gene Ontology (GO) enrichment was calculated with FUNAGE-Pro v2 (56).

### Data availability

Sequencing data have been deposited at NCBI Gene Expression Omnibus (GEO) under accession number XXXXXX

This study does not describe any novel programs, software, or algorithms.

## RESULTS

### Direct RNA sequencing (DRS) using ONT

*E. coli* MG1655 (WT) was grown to mid-exponential growth phase in LB medium at 37 °C and RNA was extracted from two independent biological replicates. Due to the high abundance of tRNA and rRNA molecules, a tRNA/rRNA depletion strategy was applied (**Figure 1A**). Specifically, tRNA was depleted using an RNA purification column, which can eliminate RNAs of < 200 nt. The RNA was then polyadenylated, and 16S and 23S rRNAs were depleted using a specific oligo-based rRNA depletion kit to generate mRNA-enriched samples (“WT mRNA”) (**Figure 1A**). At each step, the quantity and quality of RNA were assessed by RNA chip electrophoresis. tRNA depletion was evaluated based on the signal at 100 nt and rRNA depletion was assessed based on the presence of peaks at ∼1,500 nt (16S) and ∼2,900 nt (23S) (**Figure 1A**). The presence of 16S and 23S rRNA was also quantified in the WT mRNA samples via RT-qPCR; these samples contained 0.4–0.6% and 0.1% of the 16S and 23S rRNA levels, respectively, of the non-depleted samples (**Supplementary Figure 1A**).

**Figure 1.**
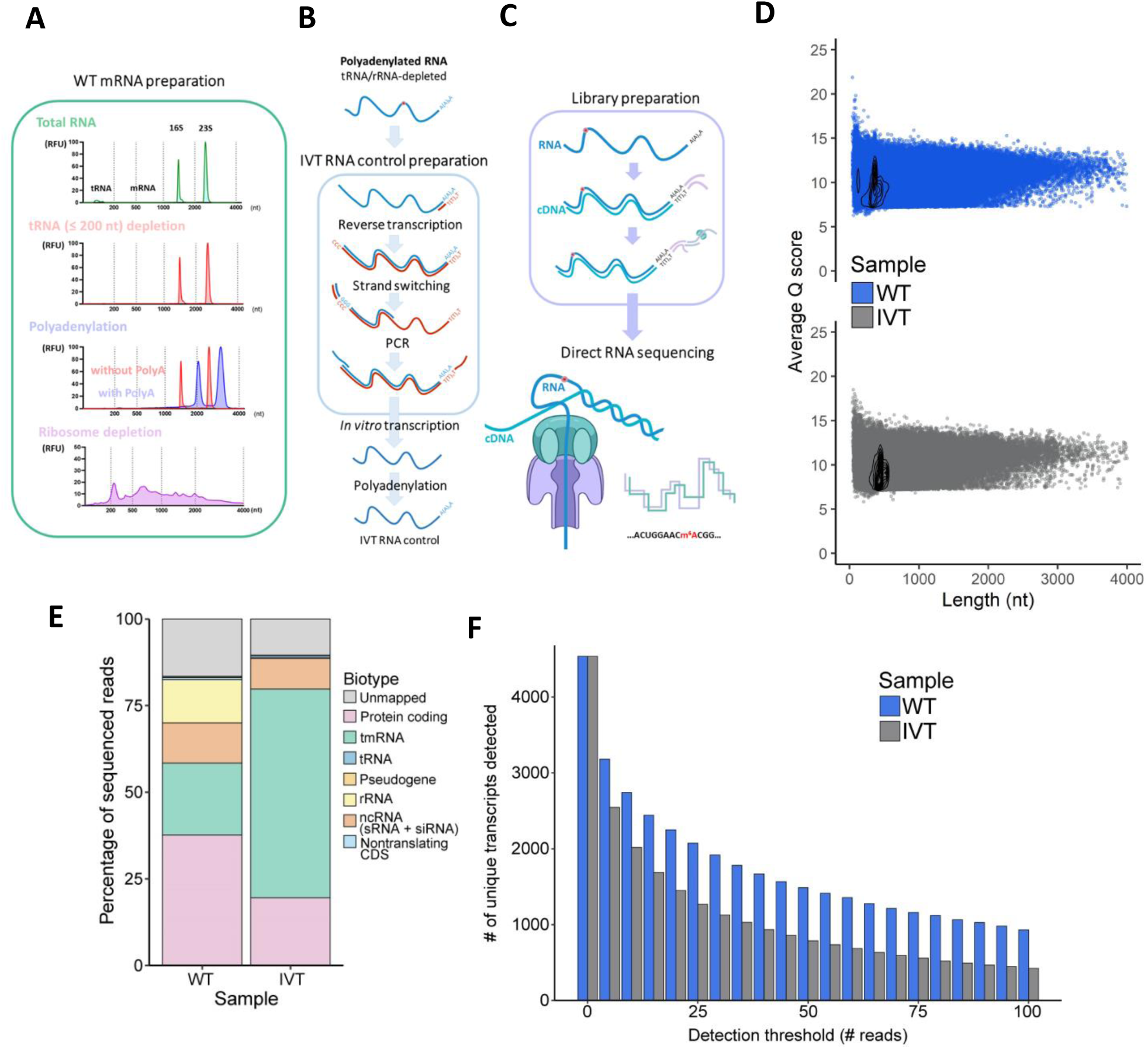
Direct RNA sequencing (DRS) of *Escherichia coli*. (A) Electropherograms from each step of enriched mRNA preparation. (B) Preparation of the *in vitro* transcribed (IVT) *E. coli* RNA control. (C) Graphical illustration of the library preparation and Nanopore sequencing protocol. Red dots represent RNA modifications. (D) Average Q score per read versus read length for the wild type (WT) (blue) and IVT (gray) RNA samples. (E) Relative proportions of bases mapping to specific RNA types in the wild-type (WT) and IVT RNA control samples. tmRNA, transfer-messenger RNA; tRNA, transfer RNA; rRNA, ribosomal RNA; ncRNA, noncoding RNA; sRNA, small RNA; siRNA, small interfering RNA; CDS, coding sequence. (F) Number of transcripts detected in the WT and IVT RNA samples across thresholds of the minimum read number.

As an unmodified RNA control, IVT RNA was prepared by adapting available protocols (42,43) for use with prokaryotic samples (**Figure 1B**). Specifically, polyadenylated (tRNA-/rRNA-depleted) RNA (**Supplementary Figure 1B**) was prepared and a single reverse primer was used to initiate reverse transcription of all molecules. A second primer was used to switch the strands and produce cDNA from *E. coli* RNA. The DNA was then amplified by PCR, incorporating the T7 promoter into the forward primer (**Figure 1B**). The resulting PCR product was used as a template for IVT, yielding unmodified RNAs (**Supplementary Figure 1C**; **Figure 1B**). After a final polyadenylation step (**Supplementary Figure 1D**; **Figure 1B**), the samples were prepared for DRS on the ONT MinION platform (**Figure 1C**). Two independent biological replicates of the WT mRNA and the IVT RNA control were sequenced individually, then the replicate samples were combined for further analyses. The combined IVT sample produced a total of ∼1.15 million reads with an average Q score of 9.2, and there were ∼1.6 million reads with an average Q score of 9.6 from the WT sample (**Table 1**; **Figure 1D**).

**Table 1.**
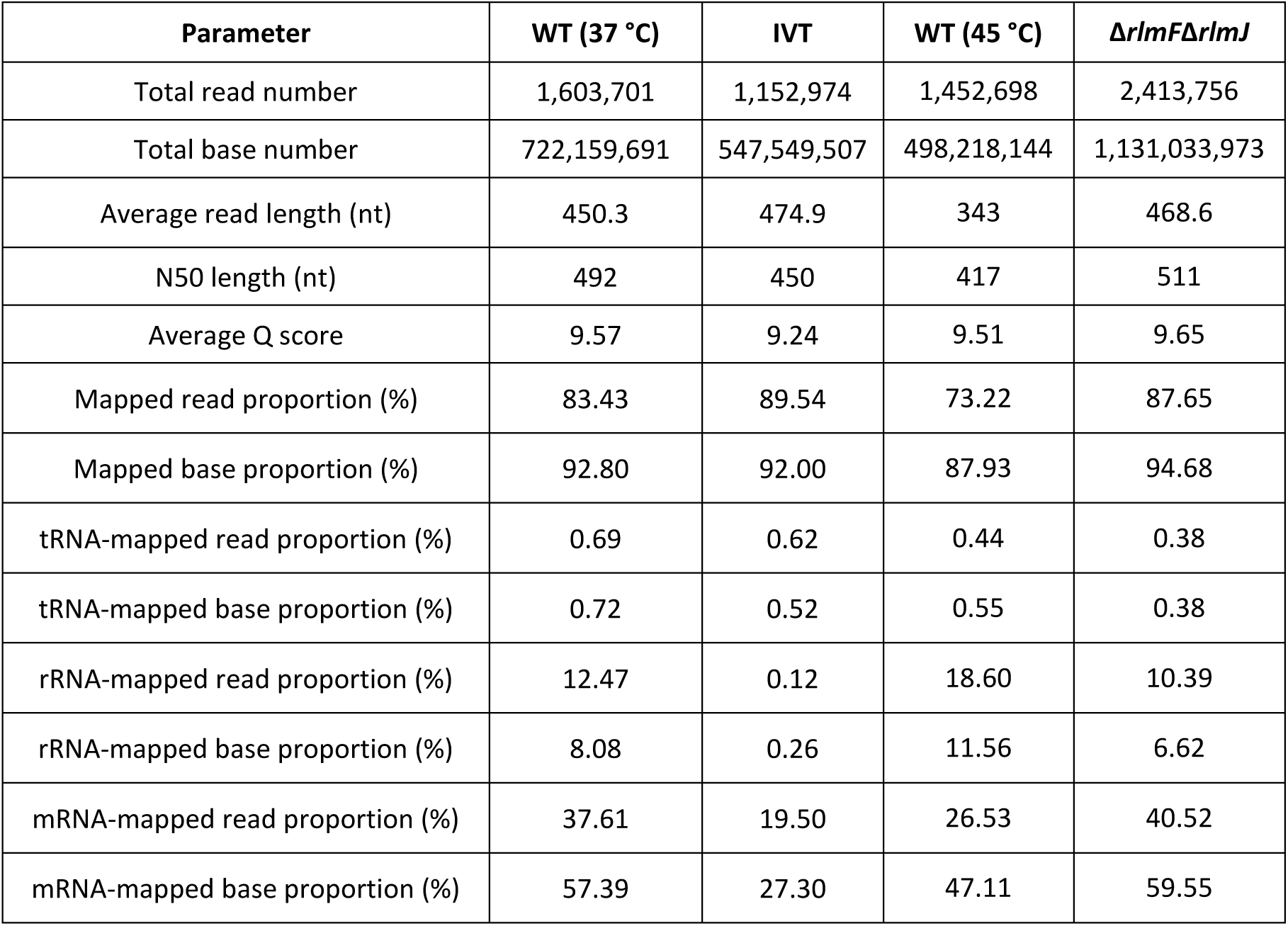
Summary of data produced with Oxford Nanopore Technologies (ONT) sequencing. Sequencing reads were generated on the ONT MinION platform. Per-sample summary statistics were produced from a combination of two biological replicates each. Heat-stressed samples are denoted “WT (45 °C)” and *E. coli* Δ*rlmF*Δ*rlmJ* double mutants are denoted “Δ*rlmF*Δ*rlmJ”*.

| Parameter | WT (37 °C) | IVT | WT (45 °C) | $\Delta rlmF\Delta rlmJ$ |
| --- | --- | --- | --- | --- |
| Total read number | 1,603,701 | 1,152,974 | 1,452,698 | 2,413,756 |
| Total base number | 722,159,691 | 547,549,507 | 498,218,144 | 1,131,033,973 |
| Average read length (nt) | 450.3 | 474.9 | 343 | 468.6 |
| N50 length (nt) | 492 | 450 | 417 | 511 |
| Average Q score | 9.57 | 9.24 | 9.51 | 9.65 |
| Mapped read proportion (%) | 83.43 | 89.54 | 73.22 | 87.65 |
| Mapped base proportion (%) | 92.80 | 92.00 | 87.93 | 94.68 |
| tRNA-mapped read proportion (%) | 0.69 | 0.62 | 0.44 | 0.38 |
| tRNA-mapped base proportion (%) | 0.72 | 0.52 | 0.55 | 0.38 |
| rRNA-mapped read proportion (%) | 12.47 | 0.12 | 18.60 | 10.39 |
| rRNA-mapped base proportion (%) | 8.08 | 0.26 | 11.56 | 6.62 |
| mRNA-mapped read proportion (%) | 37.61 | 19.50 | 26.53 | 40.52 |
| mRNA-mapped base proportion (%) | 57.39 | 27.30 | 47.11 | 59.55 |

The *E. coli* genome contains duplicate tRNA and rRNA genes encoding identical molecules that cannot be effectively distinguished from each another. Therefore, to ensure accuracy, reads were mapped to a modified version of the *E. coli* transcriptome (47) containing a single representative copy of each duplicate tRNA and rRNA gene. For example, only the 16S gene *rrsH* and the 23S gene *rrlH* were retained. Sequencing produced ∼1.3 million and ∼1.0 million reads from the WT and IVT RNA control samples, respectively, that could be mapped to this version of the *E. coli* transcriptome (**Table 1**). Per-gene sequencing read depths were quantified with Salmon (49). In the WT sample, 45% of the 83% of reads mapping to the *E. coli* genome corresponded to protein-coding regions (**Figure 1E**; **Table 1**). Approximately 2,154 transcripts (49.4% of all unique *E. coli* RNAs) were present in the WT sample at a depth of 20 or more reads (**Figure 1F**), which has been reported as the minimum depth required to accurately analyze RNA modifications (11). Of the non-mRNAs, the tmRNA *ssrA* was the most highly abundant transcript, accounting for ∼25% of all reads in the WT sample (**Figure 1E**). This was significantly more than the ∼10–15% previously detected with Illumina sequencing (57). rRNAs and tRNAs accounted for ∼15% and ∼1%, respectively, of all reads in the WT. In the IVT RNA control, 19.5% of reads mapped to mRNA sequences, corresponding to 1,379 transcripts (31.6%) that were present at a sufficient sequencing depth for further analyses (**Figure 1F**). Of the RNAs that were not present at a sufficient sequencing depth, there was no statistically significant difference in RNA size between the IVT and WT samples (**Supplementary Figure 1E**). Notably, the IVT RNA control sample had a very high level of tmRNA (∼60% of all reads), a serious distortion of the true abundance of this RNA *in vivo*. The increased abundance in the IVT samples was likely due to the presence of a poly(A) tail in the tmRNA. rRNAs and tRNAs accounted for 0.12% and 0.62%, respectively, in the IVT RNA control sample; the rRNA abundance in the final sample was dependent on the amount spiked in after IVT (see **Materials and Methods**). Even at this low read depth, at least 31 (67.4%) of the 46 unique *E. coli* tRNA molecules with known modifications were detected at a read depth of at least 20 in both the WT and IVT RNA control samples, indicating that the numerous known modifications in the tRNA and rRNA (in addition to those in other regions) could be used to validate our sequencing results.

### Detection of known RNA modifications in *E. coli*

We first searched for known modifications in the mRNA using ELIGOS (11), which identifies nt modifications by comparing the percent error of specific bases (%ESB) produced during Nanopore sequencing of a modified WT RNA sample with those in an unmodified IVT control; the proportion of %ESB in the WT compared to the IVT sample is referred to as the OddR. A recent study using Pseudo-Seq demonstrated the presence of 44 Ψ sites in *E. coli* mRNAs (18). Using ELIGOS, we were able to identify nine of these sites, which were in the CDSs of *rplA*, *tufB*, *acnB*, *rpsQ*, *tufA, secY, rpsA, ptsI*, and *fimA* (**Supplementary Figure 2A**). Moreover, the known tmRNA modifications at positions 341 (m^5^U) and 342 (Ψ) (58,59) of *ssrA* were detected in our dataset (**Supplementary Figure 2B**).

Depletion protocols are not 100% effective in removing tRNAs and rRNAs. We therefore took advantage of the presence of tRNAs and rRNAs remaining in our samples by analyzing known modifications in these reads. Importantly, the tRNA read depths were comparable between our samples and those from a prior publication describing analysis of solely tRNAs (60). Furthermore, coverage across the entire rRNA was more than sufficient for accurate detection of modifications. Comparing the WT signal to the systemic sequencing noise (as determined with the IVT RNA control), 59.7% of all known modifications in the rRNA and tRNA produced signals that were detectable with ELIGOS (**Figure 2A**). In a previous study, it was shown that the modification types m^6^A, 1-methyladenosine (m^1^A), 5-methoxyuridine (5moU), Ψ, m^7^G, inosine (Ino), 5-hydroxymethylcytidine (hm^5^C), and 5-formylcytidine (f^5^C) can be detected with ELIGOS, but m^5^C cannot (11). Similarly, we were able to detect signals for 25 of the 34 different types of modifications described in *E. coli* rRNA and tRNA, but the modification types s^4^U, 2-thiocytidine (s^2^C), m^5^C, 5-methylaminomethyluridine (mnm^5^U), 2-lysidine (k^2^C), N^6^-methyl-N^6^-threonylcarbamoyladenosine (m^6^t^6^A), I,N^6^-isopentenyladenosine (i^6^A), and N4,2′-O-dimethylcytidine (m^4^Cm) were not detected.

**Figure 2.**
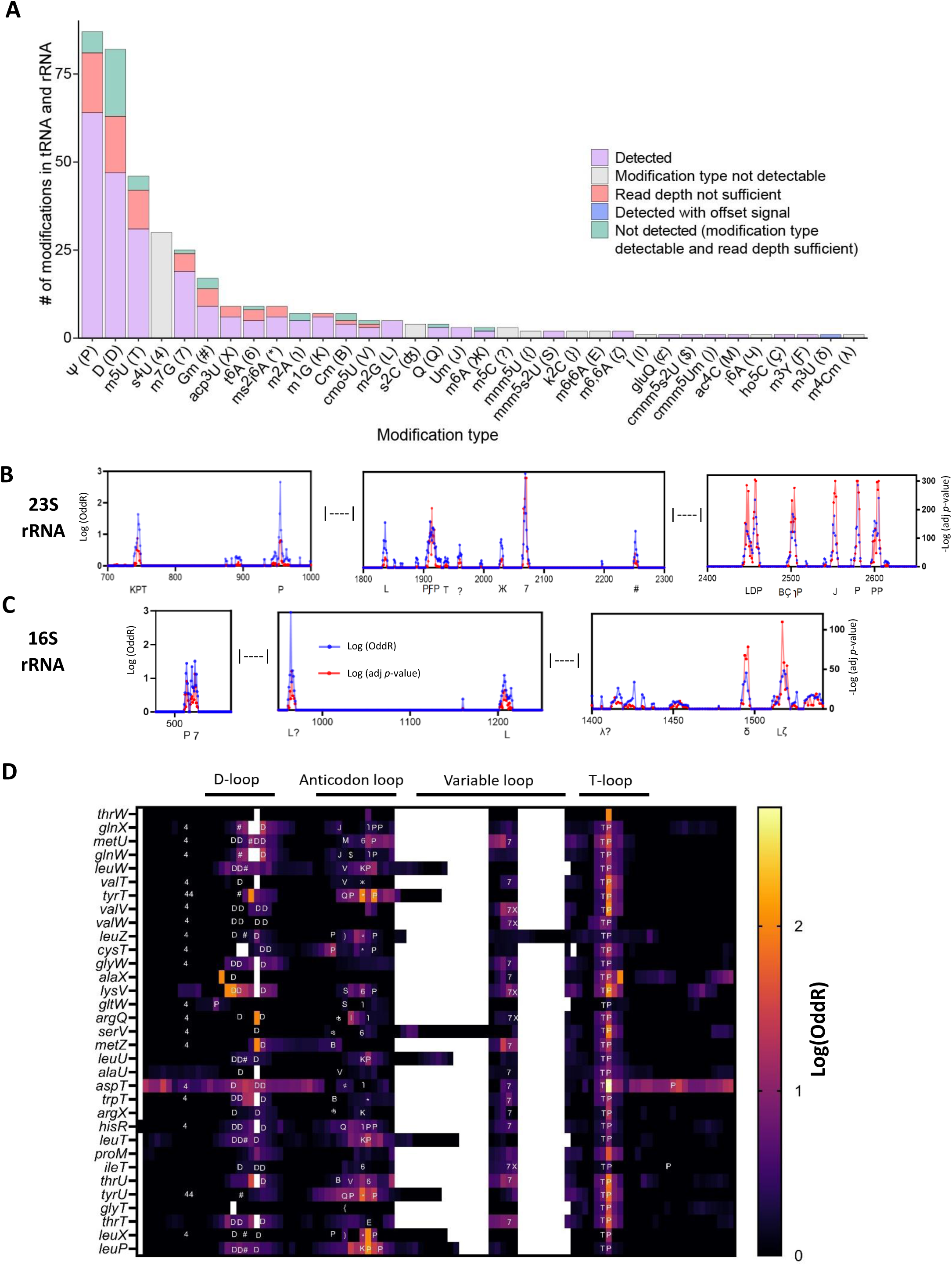
Detection of known tRNA and rRNA modifications. WT *E. coli* RNA and the IVT RNA control were processed with DRS and analyzed using ELIGOS. Putative modification sites were compared to the set of all known modifications in the *E. coli* tRNA and rRNA as published in MODOMICS [52]. (A) Overview of known tRNA and rRNA modifications. Purple indicates a modification that was detected at the known modified site; light grey indicates a modification type that is not known to be detectable with ELIGOS; pink indicates a modification in an RNA that was not present at a sufficient sequencing depth (≥ 20 reads); green indicates a modification of a type that is detectable with ELIGOS and within an RNA that was present at a sufficient sequencing depth but was not detected; and blue indicates a modification that was detected through a signal that was offset from the known modification site. (B, C) Negative log-transformed adjusted (adj) *p*-value (red) and log-transformed odds ratio (blue) values at sites across the (B) 23S and (C) 16S rRNAs. Modification types are shown below the graph. (D) Log-transformed odds ratio (OddR) values at sites across the 33 tRNAs in which modification signals were detected with ELIGOS. Symbols associated with the modification types are consistent with those used in the MODOMICS database [52]: D, dihydrouridine (D); *, 2-methylthio-N6-isopentenyladenosine (ms^2^i^6^A); ʤ, 2-thiocytidine (s^2^C); /, 2-methyladenosine (m^2^A); 4, 4-thiouridine (S^4^U); V, uridine 5-oxyacetic acid (cmo^5^U); M, N4-acetylcytidine (ac^4^C); ), 5-carboxymethylaminomethyl-2’-O-methyluridine (cmnm^5^Um); $, 5-carboxymethylaminomethyl-2-thiouridine (cmnm^5^s^2^U); S, 5-methylaminomethyl-2-thiouridine (mnm^5^s^2^U); {, 5-methylaminomethyluridine (mnm^5^U); K, 1-methylguanosine (m^1^G); 6, N6-threonylcarbamoyladenosine (t^6^A); P, pseudouridine ( Ψ); T, 5-methyluridine (m^5^U); L, N2-methylguanosine (m^2^G); Ƒ, 3-methylpseudouridine (m^3^Y); ?, 5-methylcytidine (m^5^C); E, N6-methyl-N6-threonylcarbamoyladenosine (m^6^t^6^A); Ж, N6-methyladenosine (m^6^A); 7, 7-methylguanosine (m^7^G); J, 2’-O-methyluridine (Um); #, 2’-O-methylguanosine (Gm); B, 2′O-methylcytidine (Cm); Ç, 5-hydroxycytidine (ho^5^C); ɿ, 2-methyladenosine (m^2^A); λ, N4,2’-O-dimethylcytidine (m^4^Cm); δ, 3-methyluridine (m^3^U); ζ, N6,N6-dimethyladenosine (m^6,6^A); ⊄, glutamyl-queuosine (gluQ); Q, queuosine (Q); X, 3-(3-amino-3-carboxypropyl)uridine (acp^3^U).

After controlling for sequencing noise based on the IVT sample, we identified 31 out of 36 known rRNA modifications (86.1%) in the WT mRNA sample. However, excluding the types of modifications that are not known to be detectable with ELIGOS left 32 known rRNA modifications; i.e., 96.8% of the expected modifications were detected (**Figure 2A**). m^6^A (Ж) could be detected at position 2030 but not at position 1618 of the 23S rRNA (**Figure 2B, C**), which is consistent with a previous report (17). It was not possible to identify m^5^C at C967/C1407 (L) of the 16S rRNA or C1962 of the 23S rRNA, or to detect m^4^Cm at position 1402 of the 16S rRNA (**Figure 2C**), presumably because the signal alterations produced by m^5^C and m^4^Cm were not strong enough to be classified as errors during basecalling (11). Notably, m^3^U (δ) modification of the 16S rRNA did not produce a signal at the known modification site (position 1498), but did produce an offset signal 5 nt upstream (**Figure 2A**, **C**; **Supplementary Figure 3A**). In the tRNA, we were able to detect signals in 198 out of all 346 known modification sites (57.2%) (**Figure 2D**, **Supplementary Figure 3B**). Considering only the modifications that are known to be detectable with ELIGOS and the tRNAs that were present at a sufficient sequencing depth in our dataset (72.7% of the unique tRNA molecules with known modifications), we were able to detect 83.2% of known tRNA modifications (**Supplementary Figure 3B**). Overall, these findings confirmed the validity of our approaches for RNA modification detection.

We next benchmarked the performance of Nanocompore (52) and DRUMMER (13) using the set of all known modification sites in the *E. coli* tRNA and rRNA (61). In the Nanocompore results, putative modification sites were considered only if they had a *p*-value < 0.05 from the Gaussian mixture model (GMM) logit test or from the Kolmogorov-Smirnov test for current intensity or dwell time; sites in the DRUMMER results were considered if they had an OddR ≥ 1.5 and adjusted *p*-values < 0.05 for the G-test and the OddR. In total, Nanocompore detected 24.9% of the expected tRNA and rRNA modifications (**Supplementary Figure 4A**); after exclusion of RNAs that were not present at a depth of ≥ 20 reads, Nanocompore identified 32.4% of the known tRNA and rRNA modifications. DRUMMER detected just 7.9% of the expected tRNA and rRNA modifications (**Supplementary Figure 4B**). Excluding RNAs that were not present at a sufficient read depth, this increased to 10.2%. Thus, ELIGOS showed superior performance compared to the other two programs we tested that are capable of detecting a range of modification types, and ELIGOS results were prioritized for further analyses.

### Detection of m^5^C in *E. coli* RNA

Previous studies have evaluated the presence of m^5^C in *E. coli* RNA using bisulfite treatment coupled with RT and DNA sequencing (62). Using this strategy, known m^5^C sites have been detected at positions 967 and 1407 of the 16S rRNA and 1962 of the 23S rRNA, but due to a lack of sequencing depth in the CDS, the presence of m^5^C in *E. coli* mRNA could neither be confirmed nor refuted (62). Although ELIGOS is also unable to detect m^5^C modifications, another program developed for the analysis of Nanopore data, CHEUI, does not have this limitation (51). CHEUI can be run in CHEUI-diff mode to evaluate the differences between paired samples (e.g., WT compared to IVT RNA or a mutant sample) or in CHEUI-solo mode to calculate the probability that a given 9-mer is modified. Here, a comparison of the WT mRNA to the IVT RNA control was performed with CHEUI-diff using thresholds of stoichiometry difference ≥ 0.1 and adjusted *p*-value ≤ 0.05 (**Figure 3A**). Samples were also processed individually with CHEUI-solo, considering only sites with a probability ≥ 0.9999 in the WT and < 0.999 in the IVT RNA control. Seven 9-mers were detected using this approach (**Figure 3B**): the known m^5^C modification at position 967 of the 16S rRNA; the known m^5^C modifications at site 1962 of the 23S rRNA and the adjacent C (**Figure 3A**); ho^5^C at site C2501 of the 23S rRNA; 2’-O-ribose (Cm) at site C2498 of the 23S rRNA and the adjacent C (**Figure 3A**); and site 56 in the tRNA *thrU* (genomic position 4175443). The signal in *thrU* was presumably produced by nearby known modifications, namely the previously reported modifications at the side of the T-loop (sites 54 and 55). Using less stringent thresholds, the m^4^Cm at position 1402 of the 16S rRNA could be detected with CHEUI m^5^C (stoichiometry difference 0.51, adjusted *p*-value 3.87 × 10^−32^, and probability 0.999), as could the m^5^C at site C1407 of the 16S rRNA (stoichiometry difference 0.42, adjusted *p*-value 2 × 10^−11^, and probability 0.989). The stoichiometry values predicted from Nanopore sequencing of the WT and IVT RNA control samples and the calculated stoichiometry differences were plotted for the regions surrounding known m^5^C sites in *E. coli* at positions 967 (**Figure 3C**) and 1407 (**Figure 3D**) of the 16S rRNA and 1962 of the 23S rRNA (**Figure 3E**). These well-characterized modifications could be clearly distinguished from background noise using the stoichiometry predictions generated with CHEUI. Furthermore, consistent with previous results from bisulfite sequencing (62), m^5^C methylation levels were lower at the 16S rRNA position 1407 than at the 16S position 967 or the 23S rRNA position 1962 (**Figure 3D**). This recapitulation of earlier findings demonstrates the reliability of our approach for detecting RNA modifications.

**Figure 3.**
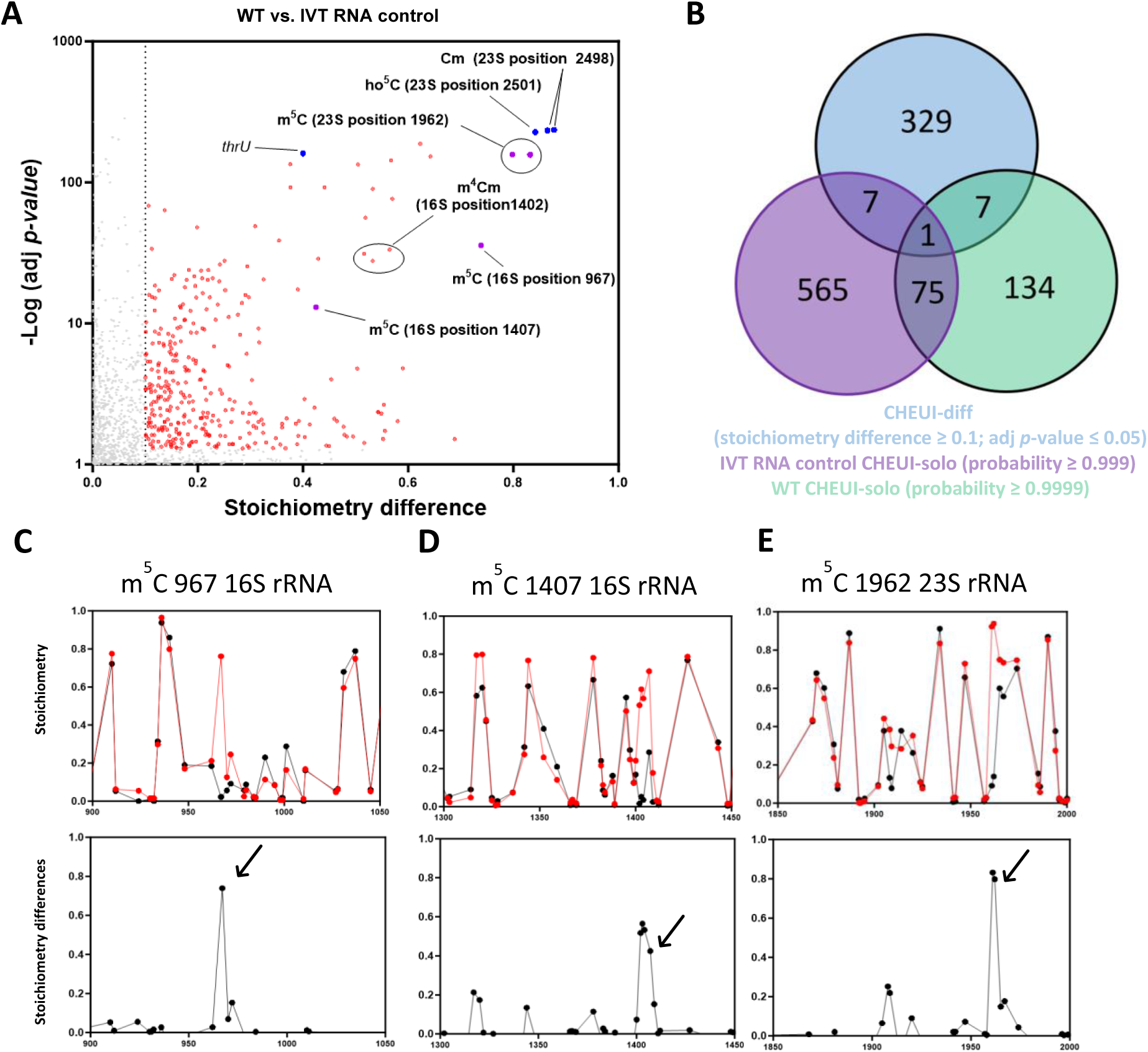
Detection of m^5^C in *E. coli* RNA. (A, B) WT *E. coli* RNA and IVT RNA control samples were processed with DRS and analyzed using CHEUI-diff for m^5^C. Each sample was also analyzed using CHEUI-solo. (A) CHEUI-diff data for m^5^C in a comparison of WT *E. coli* RNA with the IVT RNA control. Each point represents a 9-mer. Red points correspond to 9-mers with significant differences in stoichiometry at a threshold of ≥ 0.1 and adj *p* ≤ 0.05; blue points represent positions that were identified with CHEUI-solo at a probability ≥ 0.9999 in the WT sample, and a probability ≤ 0.999 in the IVT sample and with CHEUI-diff at a stoichiometry difference ≥ 0.1 and adj *p* ≤ 0.05; and purple points represent known m^5^C sites at the 967 and 1407 positions of the 16S rRNA and position 1962 of the 23S rRNA. (B) Total numbers of 9-mers identified with CHEUI-diff in the comparison of WT *E. coli* to the IVT RNA control (blue), with CHEUI-solo in the IVT RNA control (purple), and with CHEUI-solo in WT *E. coli* RNA (green). (C–E) Upper panels, CHEUI-diff stoichiometry values for m^5^C in the comparison of WT *E. coli* with IVT RNA at each 9-mer in the 16S and 23S rRNA transcripts. Lower panels, stoichiometric differences at each position in the 16S and 23S rRNA transcripts. Graphs are shown for a region of 150 nt around three known modified positions: (C) m^5^C at position 967 of the 16S rRNA, (D) m^5^C at position 1407 of the 16S rRNA, and (E) m^5^C at position 1962 of the 23S rRNA. Arrows indicate 9-mers containing known modified nucleotides.

### Detection of m^6^A in *E. coli* RNA

To evaluate m^6^A (the most prevalent mRNA modification in eukaryotes) in prokaryotic mRNA, mutants lacking the previously identified *E. coli* m^6^A rRNA methyltransferase genes *rlmF* (63) and *rlmJ* (39) were generated. We also examined the mutant *trmM,* which lacks the gene encoding the methyltransferase responsible for m^6^A methylation of the tRNA^Val^ position A37 (64). MS was used to quantify m^6^A contents in rRNA + mRNA (tRNA-depleted) samples of several *E. coli* lines: the WT, the single mutants Δ*rlmF*, Δ*rlmJ*, and Δ*trmM*, the double mutant Δ*rlmF*Δ*rlmJ*, and Δ*rlmF*Δ*rlmJ* complemented *in cis* with *rlmF* and *rlmJ*. The WT strain and the complemented mutant had similar m^6^A levels (**Figure 4A**). The single deletion mutants Δ*rlmF* and Δ*rlmJ* showed half the methylation levels of the WT strain, and m^6^A was barely detectable in the double deletion mutant (**Figure 4A**). The Δ*trmM* mutant had similar m^6^A contents as the WT, suggesting that TrmM did not play a role in rRNA or mRNA methylation. To confirm the functionality of TrmM, tRNAs (< 200 nt) were purified from WT and Δ*trmM E. coli*; the mutant showed an eight-fold decrease in tRNA m^6^A levels compared to the WT strain, confirming its role as a tRNA m^6^A methyltransferase (**Supplementary Figure 5A**) and supporting a lack of TrmM involvement in rRNA or mRNA methylation.

**Figure 4.**
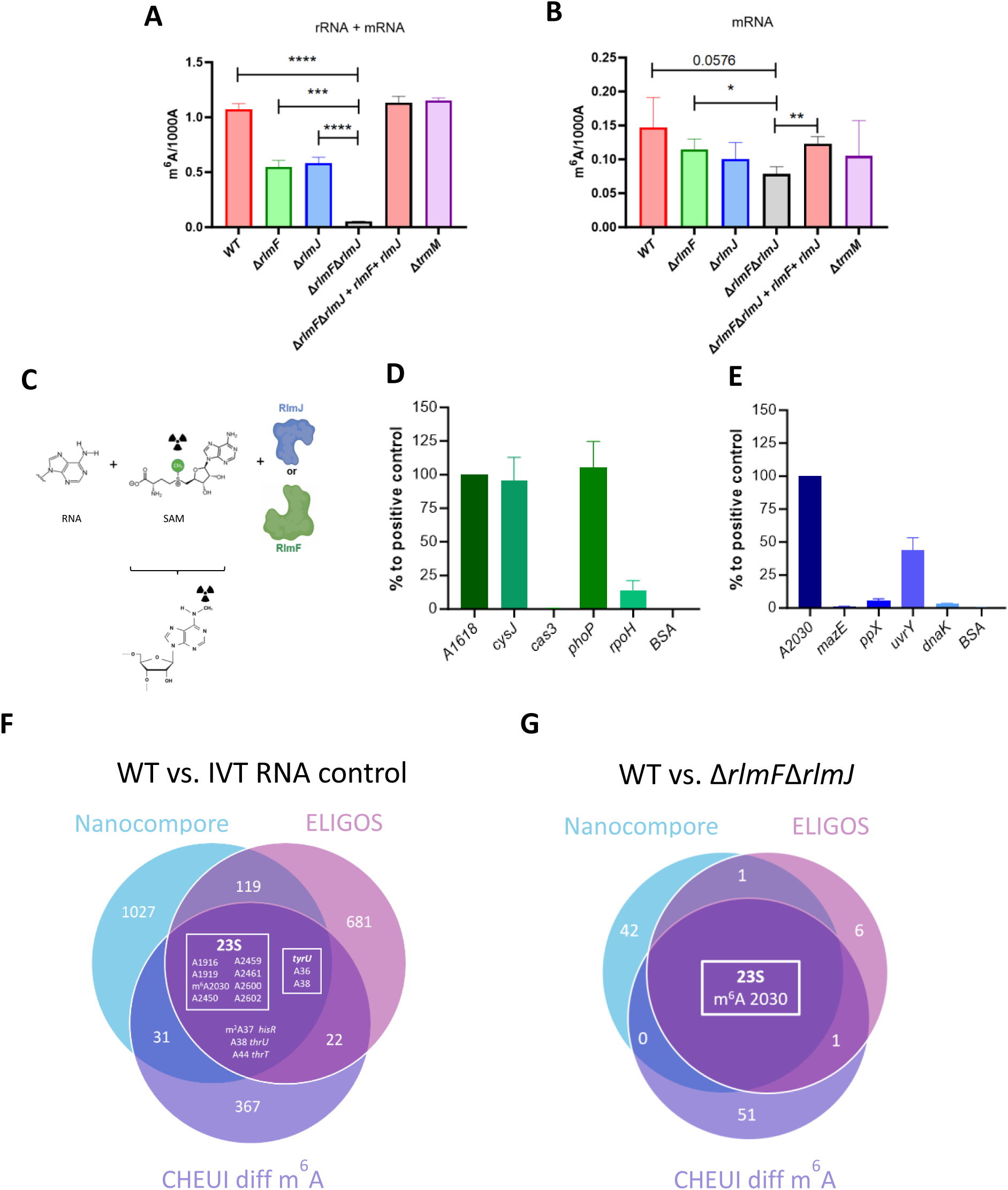
Detection of m^6^A in *E. coli*. (A, B) Total RNA samples were purified from WT *E. coli*; the single mutants Δ*rlmF,* Δ*rlmJ,* and Δ*trmM*; the Δ*rlmF*Δ*rlmJ* double mutant; and the Δ*rlmF*Δ*rlmJ + rlmF + rlmJ* complementation strain. tRNA was depleted from total RNA to obtain rRNA + mRNA samples and both tRNA and rRNA were depleted from total RNA to produce mRNA-enriched samples (mRNA). m^6^A levels were measured via mass spectrometry in (A) rRNA + mRNA (tRNA-depleted) samples and (B) mRNA-enriched (tRNA-/rRNA-depleted) samples. Data are presented as the number of m^6^As per 1000 non-modified As (1000 A). \**p* < 0.05, \*\**p* < 0.01, \*\*\**p* < 0.001, \*\*\*\**p* < 0.0001; ns, not significant (unpaired Student’s *t*-test). (C) Graphical illustration of the *in vitro* methylation protocol. (D, E) Methyltransferase activity of purified (D) RlmF and (E) RlmJ. IVT RNAs of 100–300 nt were generated for *cysJ, cas3, phoP, rpoH, mazE, ppX, uvrY*, and *dnaK*, each of which contained an expected modification site; 30-nt RNAs containing As in the appropriate sequence context were also generated for the 23S rRNA positions 2030 and 1618. (F, G) Unique and overlapping putative modification sites identified in WT *E. coli* RNA compared to RNA from (F) the IVT control and (G) Δ*rlmF*Δ*rlmJ* mutants using Nanocompore (blue), ELIGOS (pink), and CHEUI-diff for m^6^A (purple).

METTL16 is an RlmF homolog (65) that has been shown to methylate MAT2A mRNA and U6 small nuclear RNA (snRNA) in eukaryotes (66). Because transcription and translation occur in a single compartment in prokaryotes, we tested whether one of these methyltransferases could also methylate mRNA in *E. coli*. As described above, mRNA was enriched through tRNA/rRNA depletion (**Figure 1A**); rRNA was reduced to maximum levels of 1% (16S rRNA) and 0.3% (23S rRNA) of the total RNA (**Supplementary Figure 5B, C**), indicating successful mRNA enrichment. Samples from WT *E. coli*, the complemented Δ*rlmF*Δ*rlmJ* mutant, and the Δ*trmM* mutant showed comparable m^6^A levels (**Figure 4B**). In the mRNA of the Δ*rlmF*Δ*rlmJ* double deletion mutant, we detected 0.078 m^6^A per 1000 A, compared to 0.147 m^6^A per 1000 A in the WT mRNA (**Figure 4B**). This suggested that the rRNA methyltransferases RlmF and RlmJ, but not TrmM, may be partially responsible for methylation of *E. coli* mRNA *in vivo*.

Next, *in vitro* methylation assays were performed to evaluate whether these methyltransferases could methylate additional sequences beyond those previously reported in the rRNA. First, the *in vitro* activities of purified RlmF and RlmJ (**Figure 4C**) were measured using fragments of 23S rRNA containing known methylation sites as positive controls. Specifically, a previously reported 28-nt fragment containing A2030 of the 23S RNA with an additional 5′-GG sequence (39) was used to test RlmJ activity. Similarly, a 30-nt fragment from 1601 to 1631 of the *E. coli* 23S rRNA (containing the known modification site A1618) was used as the positive control for RlmF activity. After confirming that the two enzymes could methylate the positive controls (**Figure 4D, E**), we examined their activity on various *in vitro*-synthesized RNA oligos corresponding to genes containing the sequences CACAGG or GUGAAGA (the consensus sequences for RlmF and RlmJ, respectively). The results indicated that RlmF was indeed able to methylate As in sequences containing the consensus motif: *cysJ* (214 nt) and *phoP* (251 nt) were methylated at levels comparable to the positive control A1618 (**Figure 4D**). A sequence derived from *rpoH* (239 nt) was methylated, albeit at a lower level than the rRNA-derived positive control sequence (**Figure 4D**). RlmJ was able to methylate the 241-nt *uvrY-*derived sequence at intermediate levels and the *ppX* (188 nt) and *dnaK* (262 nt) sequences at a low level (**Figure 4E**). Fragments derived from *cas3* (243 nt) and *mazE* (177 nt) were not methylated, although they contained the appropriate motifs (**Figure 4D**, **E**). We also found that RlmF had greater and faster methylation activity than RlmJ (**Supplementary Figure 5D, E**). Overall, these results indicated that RlmJ and RlmF could methylate *E. coli* mRNA sequences *in vitro*, but this activity was not confirmed *in vivo*.

To detect m^6^A in *E. coli*, sequenced WT and IVT RNA control samples were compared to identify m^6^A positions throughout the *E. coli* transcriptome. Several algorithms have been developed to detect m^6^A in RNA sequenced with ONT (67). We compared three programs that are able to distinguish m^6^A modifications independent of sequence context using different features: ELIGOS, which is based on modification-induced basecalling errors (11); Nanocompore, which identifies modifications based on the electric currents generated during Nanopore sequencing (52); and CHEUI-diff m^6^A, which implements a two-stage neural network (51).

Modifications identified with the three programs were compared after applying the thresholds suggested by the authors, namely an OddR ≥ 2.5 and adjusted *p*-value ≤ 0.00001 for ELIGOS, a GMM logit *p*-value ≤ 0.01 for Nanocompore, and a stoichiometry difference ≥ 0.1 and adjusted *p*-value ≤ 0.05 for CHEUI-diff m^6^A. This yielded a total of 13 positions in the tRNA and rRNA (but none in the mRNA) that were identified by all three algorithms, referred to as shared sites. The shared sites included the known m^6^A position A2030 of the 23S rRNA (**Figure 4F**). Other shared sites included 1916 and 1919 of the 23S rRNA, which are near the known modification sites P1915 and P1917, respectively. Similarly, the shared site A2450 is close to the known m^2^G site 2445; the shared sites A2459 and A2461 are near the known modification site Ψ2457; and the shared sites A2600 and A2602 are close to the known P sites 2604 and 2605 in the 23S. All three algorithms also detected A38 in *thrU*, which is adjacent to the t^6^A modification at site 37; m^2^A37 of *hisR*; A36 and A38 of *tyrU*, which surround the ms^2^i^6^A modification at site 37; and A44 of *thrT*, which is near the m^7^G modification at site 46 (**Figure 4F**). These signals are most likely noise produced by proximity to previously identified modifications or to other modification types, such as m^2^A. To identify positions that were methylated by RlmF or RlmJ specifically, we isolated RNA from the Δ*rlmF*Δ*rlmJ* mutant, depleted the sample of tRNA and rRNA, and conducted ONT sequencing (**Supplementary Figure 5F, G**). The data were then analyzed with ELIGOS, Nanocompore, and CHEUI-diff m^6^A, comparing the Δ*rlmF*Δ*rlmJ* sample to the WT (**Figure 4G**). This showed only the presence of the known modification at position A2030 of the 23S rRNA; thus, even if RlmF and RlmJ are able to methylate mRNA, the modifications occur at very low levels that can be only detected with MS at this time.

Photo-crosslinking-assisted m^6^A-seq (PA-m^6^A-seq) has previously been conducted in *E. coli* mRNA and identified 265 m^6^A peaks corresponding to the transcripts of 213 *E. coli* genes at a resolution of ∼20– 200 nt (19). Recently, another group implemented an approach based on Nanopore sequencing and MeRIP-Seq of native bacterial RNA and modification-free IVT RNA samples (21). Considering only putative modification sites both present within MeRIP peaks and supported by at least four DRS-based modification detection tools yielded 75 high-confidence m^6^A positions across *E. coli* mRNAs corresponding to 21 genes (21). To determine whether these previously identified positions could be validated using orthogonal (i.e., non-sequencing-based) methods, we performed experimental validation with the single-base elongation- and ligation-based qPCR amplification method (SELECT) (40). This qPCR-based detection approach relies on the fact that m^6^A hinders the single-base elongation activity of BstII DNA polymerase and the nick ligation efficiency of SplintR ligase. The threshold cycle (C_T_) values can then be compared between pairs of oligos, with one targeting a suspected m^6^A site and the other targeting a nearby unmodified A as a control (**Figure 5A**). To account for possible differences in annealing efficiencies between oligos, IVT RNA was employed as a negative control. The presence of an m^6^A modification at the target site would delay amplification, increasing the difference in C_T_ values (ΔC_T_) between the target site and the adenosine control (A control). As an additional control for the known m^6^A positions A1618 and A2030 in the *E. coli* 23S rRNA, RNA extracted from the double deletion mutant Δ*rlmF*Δ*rlmJ,* which lacks the *E. coli* m^6^A rRNA methyltransferases that methylate these sites (39,63), was also tested. For each suspected m^6^A site, ΔC_T_ was calculated for the WT RNA sample (ΔC_T (WT)_) and the IVT (ΔC_T (IVT)_). For the known m^6^A sites in the 23S, we also calculated ΔC_T_ for Δ*rlmF*Δ*rlmJ* RNA (ΔC_T (Mut)_). Positions were considered modified only if ΔC_T (WT)_ – ΔC_T(IVT)_ (or ΔC_T (WT)_ – ΔC_T(Mut)_) was ≤ -1 (41).

**Figure 5.**
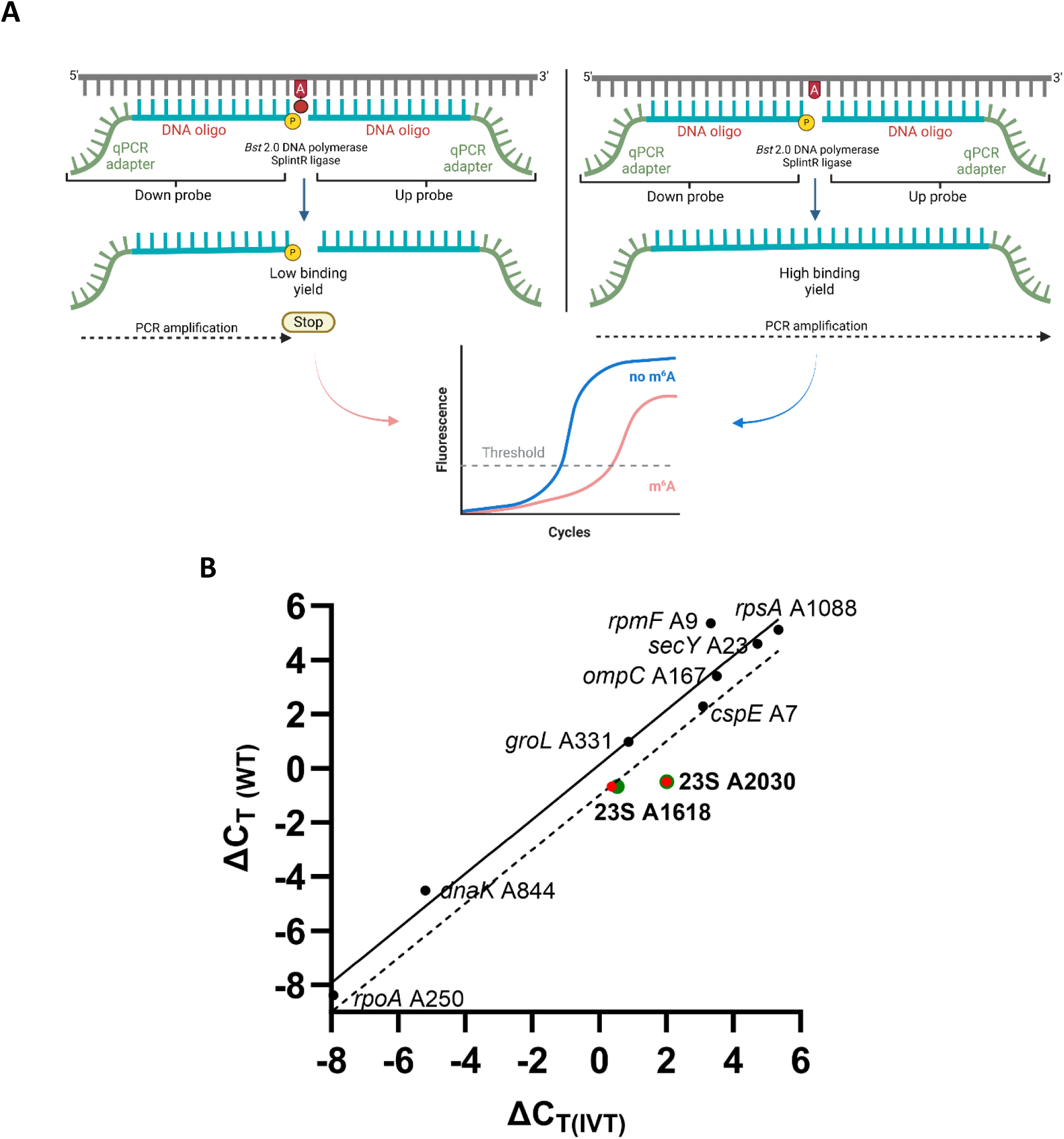
SELECT analysis of high-confidence m^6^A sites in *E. coli* mRNA. (A) Graphical illustration of the SELECT protocol. (B) Differences in the detection thresholds between several positions where m^6^A modifications were expected and nearby unmodified As (ΔC_T_) in the IVT RNA control (ΔC_T(IVT)_) and WT RNA (ΔC_T (WT)_). The solid diagonal line represents a simple linear regression of the data obtained from the putative m^6^A positions; the dotted line represents the same regression but with a difference of -1 ΔC_T_ between the WT and IVT samples. Modified positions appear below the dotted line. For positions 1618 and 2030 of the 23S rRNA, values for both the comparison of the WT with the IVT control (red) and with RNA extracted from the Δ*rlmF*Δ*rlmJ* double mutant (green) are shown.

For position A1618 of the 23S, ΔC_T (WT)_ - ΔC_T(IVT)_ was -1.03 and ΔC_T (WT)_ – ΔC_T(Mut)_ was -1.17; at position A2030, both ΔC_T (WT)_ - ΔC_T(IVT)_ and ΔC_T (WT)_ – ΔC_T(Mut)_ were -2.5 (**Figure 5B**, **Supplementary Figure 6A**). This confirmed that SELECT could be used as an orthogonal method to evaluate the presence of m^6^A in *E. coli* RNA samples. Next, we used SELECT to evaluate eight sites within eight different mRNAs that were recently identified as high-confidence *E. coli* mRNA m^6^A modifications (*secY* A23, *cspE* A7, *ompC* A167, *dnak* A844, *groL* A331, *rpsA* A1088, *rpoA* A250 and *rpmF* A9) (21). The C_T (WT)_ – ΔC_T(IVT)_ values ranged from 2.03 (*rpmF* position A9) to -0.82 (*cspE* position A7) (**Figure 5B**, **Supplementary Figure 6B**), which were above the threshold for classification as modified sites (≤ -1). Thus, SELECT did not support the presence of these m^6^A modifications in *E. coli* mRNA.

### Effects of heat stress on rRNA modifications in *E. coli*

After establishing several techniques to study the epitranscriptome of *E. coli*, we focused on assessing stress-dependent epitranscriptomic changes, specifically during heat stress. Most bacteria constantly monitor temperature changes, and the expression of many virulence genes is coordinated in response to changing temperatures (68). Sensing and responding to changes in temperature occurs particularly at the level of translation via RNA thermometers. RNA thermometers are complex RNA structures that change conformation in response to differences in temperature (69). We here asked whether changes in the epitranscriptome represent another level of bacterial regulation under heat stress.

*E. coli* cells pre-grown in LB medium at 37 °C to an OD_600_ of 0.5 were heat-stressed by the addition of hot LB to instantaneously raise the temperature to 45 °C, followed by incubation at 45 °C for 30 min to an OD_600_ of ∼1 (**Figure 6A**). The previously described *E. coli* samples grown at 37 °C to a similar OD_600_ were used as controls (**Figure 1D**). Total RNA was extracted from the cells, then tRNA and rRNA were depleted and the resulting mRNA-enriched samples were sequenced using ONT as described above. Overexpression of *dnaK,* which encodes the chaperons heat shock protein 70 (70), is often used as an indicator for the heat shock response in *E. coli*. RT-qPCR demonstrated 5.6× upregulation of *dnaK* in the heat-treated sample compared to the control sample (**Figure 6B**). Furthermore, analysis of DRS data derived from the 45 °C compared to the 37 °C samples showed upregulation of 15 genes that are regulated by the heat-shock sigma factor sigma 32 (24) (**Figure 6C**). Taken together, these results confirm that the increase in temperature triggered the heat-shock response.

**Figure 6.**
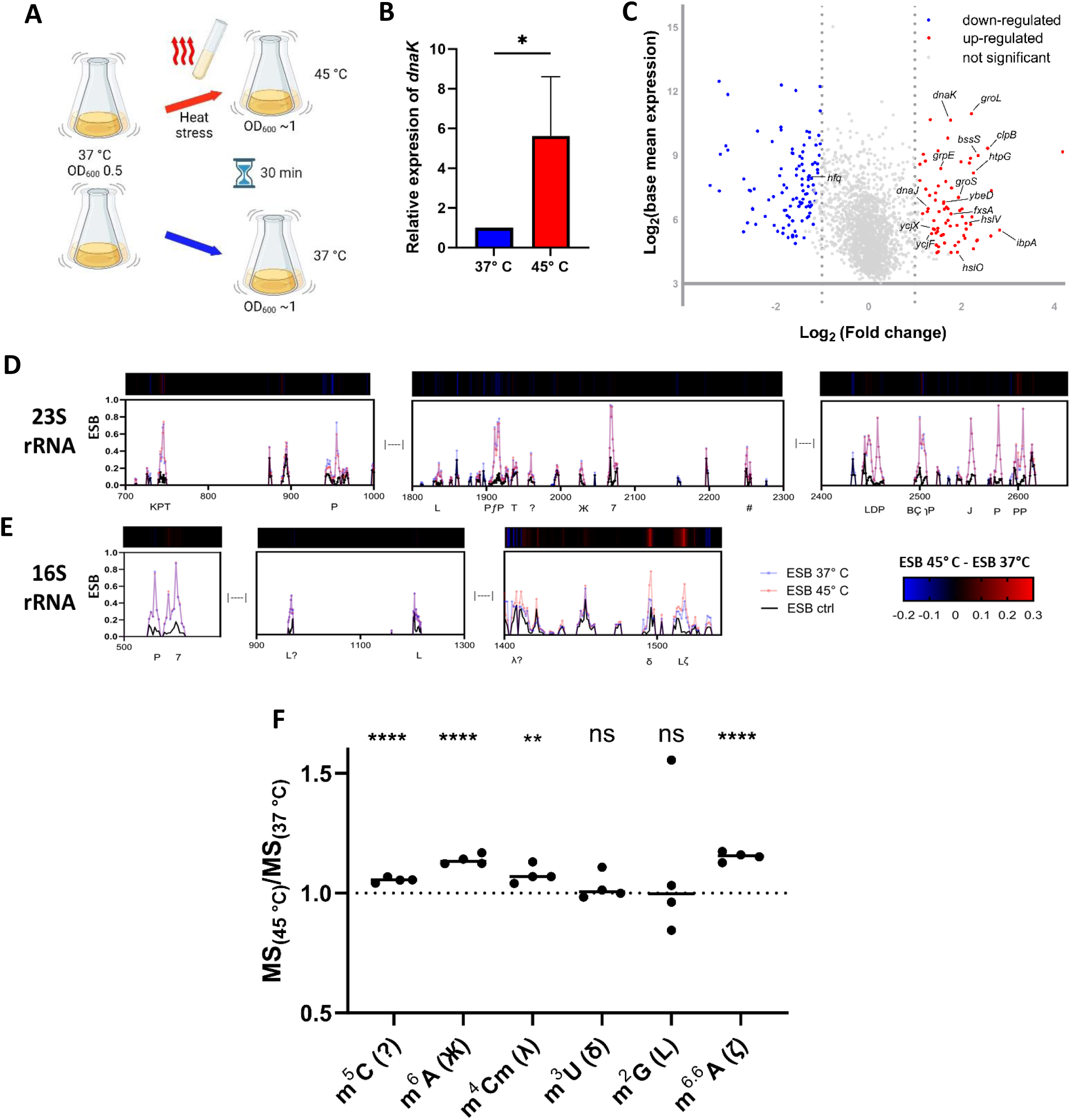
Effects of heat stress on the *E. coli* epitranscriptome. (A) Overview of the heat stress protocol. Two flasks of *E. coli* were grown in parallel at 37 °C to an OD_600_ of 0.5. LB heated to 90 °C was added to one flask to raise the temperature to 45°C and cultivation was continued at 45 °C for 30 min. The other flask was left at 37 °C. Cells were then harvested and RNA was extracted. (B) Reverse transcription quantitative PCR (RT-qPCR) was performed to evaluate expression of *dnaK*, which encodes the 70 kDa heat shock protein. *dnaK* expression was normalized to the 16S rRNA as an internal control. (C) Differentially expressed genes in *E. coli* at 45 °C compared to 37 °C as determined with DRS. Data are displayed as log-transformed base mean expression vs. log-transformed fold change. Differentially expressed genes controlled by the sigma 32 (σ^32^) factor are labeled. (D, E) ESB_test 45 °C_ (red), ESB_test 37 °C_ (blue), and ESB control (black) values for sites across the (D) 23S and (E) 16S rRNAs. The bar above each graph shows ΔESB 45 °C - Δ ESB 37 °C. Modification types are shown below each graph. (F) MS-based quantification of selected modifications in rRNA + mRNA (tRNA-depleted) samples. Values were calculated as modification abundance at 45 °C divided by abundance at 37 °C (MS_(45 °C)_/MS_(37 °C)_). \*\**p* < 0.01, **** *p* < 0.0001; ns, not significant (unpaired Student’s *t*-test).

The DRS data showed that the average read quality in the 45 °C sample (**Supplementary Figure 7A**) was comparable to that of the 37 °C sample (**Figure 1D**), as was the distribution of mapped reads (**Supplementary Figure 7B, Figure 1E**). This ensured the validity of comparisons between DRS data generated from samples grown at the two temperatures (**Supplementary Figure 7C, D**). The ESB values produced by ELIGOS can be directly used as a measure of some modification levels (11,17). We therefore compared the ESB test values of modification sites shared by the 45 °C and 37 °C samples to evaluate changes in RNA modification levels due to heat stress. There were no differences in ESB test values between the 45 °C and 37 °C samples in the 23S rRNA sites (**Figure 6D**), but temperature-specific differences were observed in the 16S rRNA. The m^6,6^A modifications at sites 1518 and 1519 (genomic positions 225,288 and 225,289, respectively) were increased in abundance at 45 °C, with ESB test values of 0.61 and 0.3, respectively, compared to 0.4 and 0.2, respectively, at 37 °C (**Figure 6E**). These observations were validated via MS to assess levels of m^6,6^A. In an rRNA + mRNA sample (tRNA-depleted), m^6,6^A was found to increase by 15% at 45 °C (**Figure 6F**). As positive controls, MS measurements were also performed for the modification m^2^G (which is present at sites G966 and G1516 of the 16S rRNA and G1835 and G2445 of the 23S rRNA); consistent with the predictions of ELIGOS, these sites showed no change in modification abundance in rRNA + mRNA samples. ELIGOS cannot detect m^5^C, (whereas CHEUI can), could not be used to assess stoichiometric changes (such as those at positions 967 and 1407 of the 16S rRNA and 1962 of the 23S rRNA), could not identify m^6^A at position 1618 of the 23S rRNA, and produced an offset signal for the m^3^U at position 1498 of 16S rRNA. We therefore assessed modification abundance at these sites with MS. Levels of m^6^A and m^4^Cm increased under heat stress (**Figure 6F**), as did m^4^Cm and m^5^C. In contrast, no changes were detected in levels of m^3^U (**Figure 6F**).

### Effects of heat stress on tRNA modifications in *E. coli*

tRNAs are highly modified molecules with known modifications clustered in the anticodon-, variable-, D-, and T-loops (**Figure 7A**). Using the tRNA modifications present in the MODOMICS database (61), which contains only thoroughly characterized, experimentally validated modifications, we examined a set of 31 tRNAs with known modifications. The ESB test value derived from ELIGOS for each modification site was used as a proxy of modification abundance (17); differences in modification levels between the 45 °C and 37 °C data were calculated from the ΔESB (ESB test – ESB control) values for both conditions.

**Figure 7.**
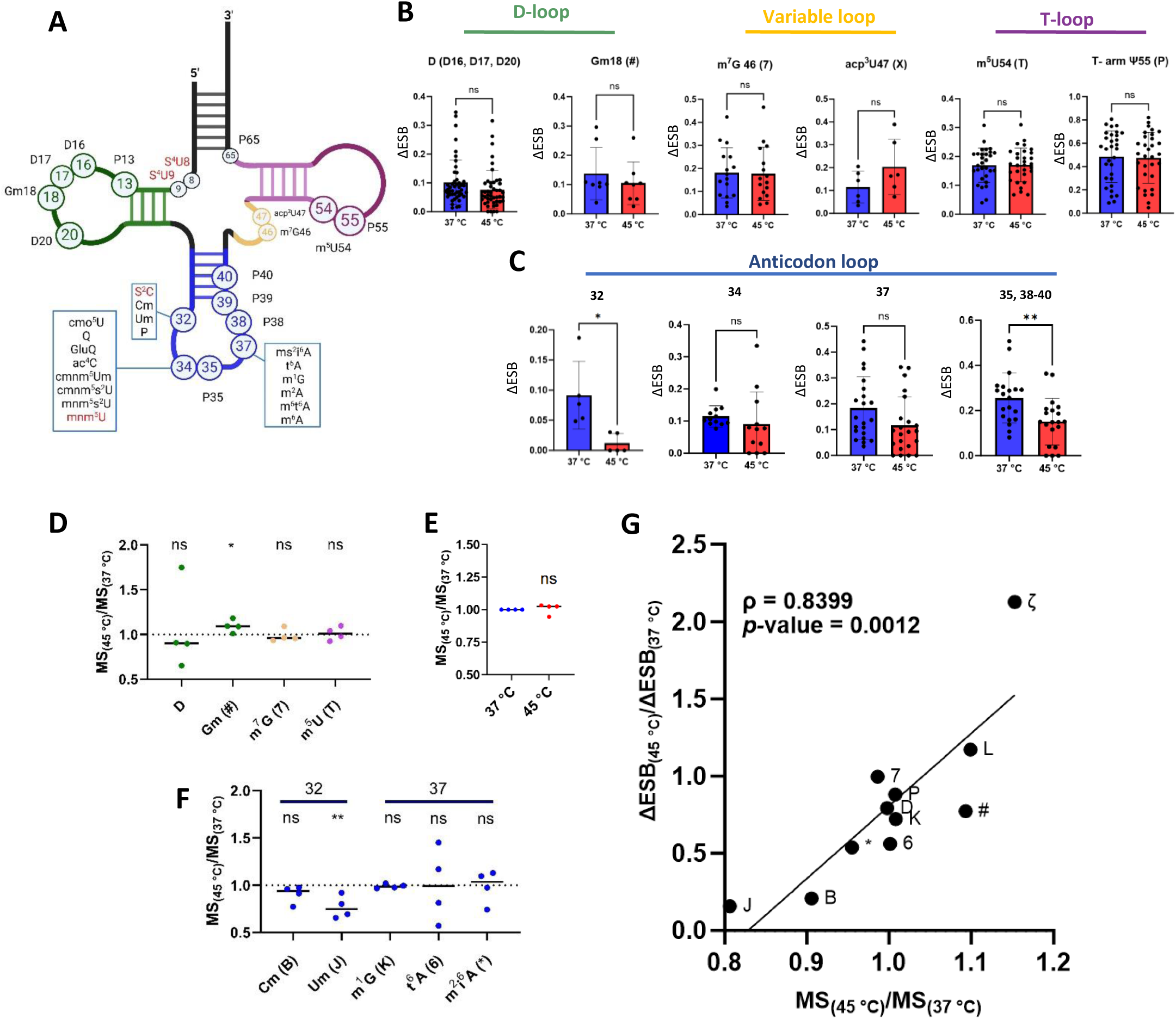
Changes in *E. coli* tRNA modifications under heat stress. (A) Schematic representation of the tRNA secondary structure and location of known modifications. Modifications marked in red were not detected in our dataset. (B, C) ΔESB values for known tRNA modification sites at 37 °C and 45 °C. ΔESB was calculated as ESB test - ESB ctrl for each condition; it was set to 0 for positions not detected in both datasets. Values are shown separately for modifications in the (B) D-, variable, T-, and (C) anticodon loops. Modifications in the D-loop comprise Gm at position 18 and dihydrouridine (D) at positions 16, 17, and 20; those in the variable loop are m^7^G (7) at position 46 and acp^3^U (X) at position 47; those in the T-loop are m^5^U (T) at position 54 and P at position 55; and those in the anticodon loop are Cm (B) and Um (J) at position 32, cmo^5^U (V), Q, GluQ (⊄), ac^4^C (M), cmnm^5^Um ()), cmnm^5^s^2^U ($), and mnm^5^s^2^U (S) at position 34, m^2^A (ɿ), ms^2^i^6^A (*), t^6^A (6), m^1^G (K), m^6^t^6^A (E), and m^6^A (Ж) at position 37, and P at positions 35 and 38–40. \**p* < 0.05, \*\**p* < 0.01; ns, not significant (unpaired Student’s *t*-test). (D–F) MS data for selected modifications in tRNA-enriched samples (RNA ≤ 200 nt). Values were calculated as modification abundance at 45 °C divided by abundance at 37 °C (MS_(45 °C)_/MS_(37 °C)_). Abundance levels are shown for (D) modifications in the D-, variable, and T-loops, (E) pseudouridine, and (F) other modifications at position 32 and 37 in the anticodon loop. \**p* < 0.05, \*\**p* < 0.01; ns, not significant (unpaired Student’s *t*-test). (G) Relative changes in modification abundance at 45 °C compared to 37 °C as measured with ELIGOS from Nanopore sequencing data (ΔESB_(45 °C)_/ΔESB_(37 °C)_) and MS (MS_(45 °C)_/MS_(37 °C)_). The correlation between modification abundance based on ELIGOS and MS data was calculated as Pearson’s correlation coefficient (ρ).

Using the ΔESB values as a measure of differences in modification abundance, we found no changes in modification levels in response to heat shock at sites in the D-loop (namely D16, D17, D20, and Gm18) (**Figure 7B**, **Supplementary Figure 8**). Similar results were found for levels of m^7^G46 and 3-(3-amino-3-carboxypropyl)uridine (acp^3^U) at position 47 in the variable loop (**Figure 7B**) and for m^5^U54 (T) and P55 levels in the T-loop (**Figure 7B**, **Supplementary Figure 8**). Several previously described modifications were evaluated by MS of tRNA-enriched samples (RNAs ≤ 200 nt) to confirm the findings of the Nanopore sequencing (**Figure 7D–F**). The results were consistent between the sequencing and MS datasets for the modification types D, m^7^G, and m^5^U (**Figure 7D**). However, an increase of ∼10% in Gm levels (as detected with MS) was not detected with sequencing (**Figure 7D**). There are multiple Ψ modifications in the tRNA T-loop and anticodon-loop; MS could be used to detect the total level of Ψ in these regions without the site specificity afforded by sequencing (**Figure 7E, Supplementary Figure 8**). Considering all of these sites together, MS showed no changes in Ψ levels at 45 °C compared to 37 °C, consistent with the sequencing results. However, sequencing did reveal significantly lower Ψ levels specifically at sites 35 and 38–40 of the anticodon at 45 °C compared to 37 °C (**Figure 7C**), highlighting the increased precision enabled by DRS. A similar decreasing trend was observed at anticodon position 32 (**Figure 7C**), which may bear one of three modification types: Cm (B), Um (J), or s^2^C (ʤ). Although s^2^C cannot currently be detected with ONT sequencing, levels of both Cm and Um at this position were found to decrease by ∼10% and 24%, respectively, at 45 °C by MS (**Figure 7F**). In contrast to these sites in the anticodon loop, modification levels at tRNA positions 34 and 37 were not found to change in response to heat stress with either ELIGOS or MS (**Figure 7C, F**). Finally, to evaluate the suitability of the ΔESB values produced by ELIGOS as a proxy for changes in modification levels, we assessed the relationship between the ratio of ΔESB_(45 °C)_ to ΔESB_(37 °C)_ and the relative changes in the MS data (MS_(45 °C)_/MS_(37 °C)_). This revealed a very high correlation rate between the datasets (ρ = 0.8399; *p* = 0.0012), supporting our use of DRS data to quantitatively evaluate changes in modification levels (**Figure 7G**). Overall, the DRS data not only corroborate the MS findings, but also offer site-level resolution of epitranscriptomic changes. This allowed, for instance, the detection of reduced D levels at specific tRNA sites, even when the averaged data (from either ELIGOS or MS) showed no changes.

### Effects of heat stress on modifications to the transcriptome in *E. coli*

To minimize errors and maximize the detection of truly modified sites with ONT sequencing, we analyzed precision and recall among tRNA and rRNA regions using thresholds of varying stringency for several parameters output by ELIGOS. Specifically, we tested a range of thresholds for the ESB test (**Supplementary Figure 9A–**), ESB control (**Supplementary Figure 9D–**), OddR (**Supplementary Figure 9G–**), and adjusted *p*-value (**Supplementary Figure 9J–**). Precision was calculated as the percentage of truly modified positions out of the total number of positions classified as modified, and recall was calculated as the percentage of known modified sites detected in our dataset out of the total number of known modified sites (see **Materials and Methods**). In addition to testing various thresholds for each parameter individually, we analyzed precision and recall for all possible combinations of these four parameters, with a total of 32,000 combinations (**Figure 8A**). Thresholding the ELIGOS results using the optimal values identified with this method (%ESB test ≥ 0.1, %ESB control ≤ 1, OddR ≥ 3, and adjusted *p* ≤ 0.05) yielded a precision of 85.9% (i.e., an error rate of 14.1%) and a recall of 58.8% in the combined tRNA/rRNA dataset.

**Figure 8.**
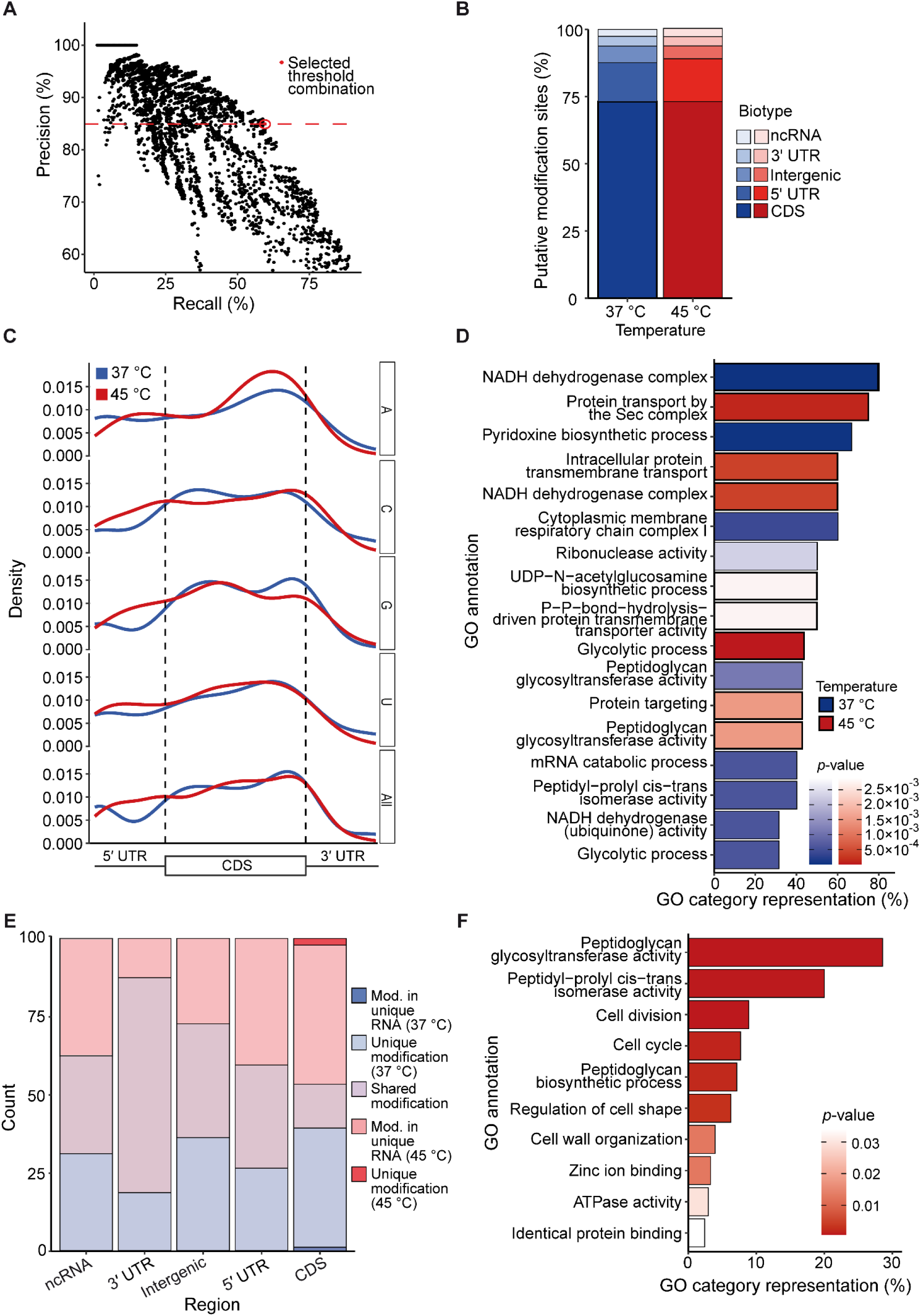
Characteristics and distribution of putative novel modification sites throughout the *E. coli* transcriptome. (A) Precision and recall of known tRNA and rRNA modification sites using 32,000 combinations of four parameters: ESB test, ESB control, odds ratio, and adjusted p-value. The combination of thresholds selected for use in further analyses is indicated with a red point. (B) Relative proportions of putative novel *E. coli* modification sites by RNA biotype, excluding tRNA and rRNA. A total of 387 and 439 sites/regions were detected outside the tRNA and rRNA in samples grown at 37 °C and 45 °C, respectively. (C) Normalized distribution of putative mRNA modifications across the 5′ untranslated region (UTR), coding sequence (CDS), and 3′ UTR in samples grown at 37 °C (blue) and 45 °C (red). (D) Gene Ontology (GO) enrichment analysis of all genes with putative mRNA modifications in samples grown at 37 °C (blue) or 45 °C (red). Adjusted *p*-values were derived from a hypergeometric probability test followed by false discovery rate (FDR) correction for multiple testing. (E) Proportions of putative modification sites shared by the 37 °C and 45 °C samples. Modifications were designated as shared, unique at 37 °C or 45 °C (unique modification), or within an RNA that was only present at 37 °C or 45 °C (modification in unique RNA). (F) GO enrichment analysis of RNAs containing putative modifications that increased in abundance at 45 °C compared to 37 °C.

These high-confidence thresholds were then applied to the remaining RNA biotypes: mRNAs, including the 5′ UTRs, CDSs, and 3′ UTRs; ncRNAs; and intergenic regions. Due to the previously demonstrated effects of modifications on neighboring nt in Nanopore sequencing (11), the signals surrounding known modifications in the tRNA and rRNA were analyzed. We consistently found detectable signals up to 10 nt upstream and downstream of known modification sites (**Supplementary Figure 10A–**). This made it difficult to determine whether the signal at each individual nt, especially in regions without well-characterized modification sites, originated from one or more closely placed modifications. Therefore, regions of adjacent nt in which all sites were classified as modified by ELIGOS were each considered a single “modification region”. After thresholding with the values established using the tRNA and rRNA data, the 37 °C dataset contained 283 putative novel modification sites/regions in the CDS (73.1% of the total), 56 in the 5′ UTR (14.5%), 24 in the intergenic region (6.2%), 14 in the 3′ UTR (3.6%), and 10 in ncRNAs (2.6%); in the 45 °C dataset, there were 321 putative modification sites or regions in the CDS (73.1%), 70 in the 5′ UTR (15.9%), 21 in the intergenic region (4.8%), 15 in the 3′ UTR (3.4%), and 12 in ncRNAs (3.1%) (**Figure 8B**).

To determine the general distribution of modifications across each transcript, the putative modification sites/regions were plotted based on their relative locations within each mRNA. This revealed enrichment of putative RNA modifications at the beginning and end of the CDS at 37 °C and in the middle and end of the CDS at 45 °C (**Figure 8C**). Nucleotide-specific analysis of the modification distributions showed that As were the most abundantly modified base in the 5′ UTR. At both 37 °C and 45 °C, modified As and Us were enriched toward the end of the CDS, whereas modified Cs and Gs showed a more bimodal distribution (**Figure 8C**). GO enrichment analysis was then performed for the putative modifications identified in mRNAs. Significantly enriched GO terms among transcripts with putative modifications at 37 °C included those related to functions in general metabolism and RNA metabolism (**Figure 8D**). At 45 °C, CDSs containing putative modifications were enriched in functions associated with transporters and cell wall synthesis (**Figure 8D**).

Comparison of the putative mRNA modification sites between the two temperature conditions revealed extensive overlap, particularly in the 3′ UTR (**Figure 8E**). Only 18 of the putative modification sites were uniquely found at 37 °C or 45 °C because the transcript containing the modification was too lowly abundant in either condition; all of the putative modification sites outside the CDS and 96.6% of those in the CDS were present in transcripts with ≥ 20 reads under both conditions, enabling direct comparisons of 133 putative mRNA modification sites/regions. As discussed above, the ESB test values produced by ELIGOS can be used as a measure of modification abundance. Thus, we compared ESB test values for putative CDS modification sites/regions present at both temperatures. Of these 74 putative modification sites/regions, a majority (59.5%) showed increased abundance at 45 °C compared to 37 °C. GO enrichment analysis was performed for transcripts containing these putative modifications that increased at 45 °C, revealing enrichment of genes related to cell wall structure and other well-characterized components of the heat stress response (**Figure 8F**). These findings suggest that mRNA modifications represent an additional layer of post-transcriptional regulation in *E. coli* under heat stress.

## DISCUSSION

Numerous epitranscriptomic studies have been conducted in eukaryotes from yeast to humans, and the machineries responsible for m^6^A and m^5^C modifications are well characterized. In comparison, mRNA modifications in bacteria remain poorly understood. One of the main reasons for this gap in epitranscriptomic knowledge is the difficulty in isolating mRNA from bacteria. The absence of poly(A) tails and the high abundance of rRNA make the mRNA isolation process a challenging endeavor. Currently, the most widely used protocols for ribosome depletion use specific biotinylated probes to eliminate rRNA. Using biotinylated probes, we were here able to reduce the 16S and 23S rRNA levels to 1% and 0.3%, respectively, of the total RNA. Although mRNA isolation methods have been greatly improved in recent years, complete rRNA removal is still not achievable (71). We used this disadvantage as an advantage, because by polyadenylating bacterial RNA prior to sequencing, we were able to analyze all RNA types (including mRNAs, ncRNAs, tRNAs and rRNAs) at the same time. Third-generation sequencing approaches provide key advantages over earlier methods: they are independent of rRNA depletion and facilitate the sequencing of native RNA and thus, the detection of modifications with single-molecule resolution without the need for potentially harmful chemical treatments. However, DRS data are extremely noisy, necessitating the analysis of unmodified controls (e.g., IVT *E. coli* RNA) to eliminate systemic error signals associated with the sequencing process itself. In addition to an IVT control, to minimize errors while maximizing the detection of truly modified sites, we used the set of known tRNA and rRNA modifications to establish an optimal combination of thresholds for several parameters associated with modification abundance and confidence. Overall, this approach proved suitable for detecting about 60% of all known tRNA and rRNA modifications in *E. coli*. Of the 34 different types of modifications known in *E. coli* tRNA and rRNA, we were able to detect 24 (including D, Ψ, m^5^U, m^7^G, and Gm) using ELIGOS (**Figure 2A**). m^5^C modifications were not detectable using ELIGOS (**Figure 2A**), but were detected using another program, CHEUI (**Figure 3A**).

Without a separate treatment such as bisulfite (62), the dataset generated in the present study was suitable for the assessment of m^5^C abundance in bacterial RNA. Although we successfully detected known m^5^C positions in addition to Cm, ho^5^C, and m^4^Cm modifications in the 16S and 23S rRNAs, the presence of m^5^C in *E. coli* mRNA was not supported. However, despite the advances of our approach compared to previous methods, the sequencing depth was not sufficient to evaluate all coding genes in the *E. coli* genome. Therefore, our results do not conclusively prove the absence of m^5^C in *E. coli* mRNA.

Using orthogonal methods, we were unable to detect previously reported methylation sites identified with PA-m^6^A-seq, a sequencing method that relies on antibody recognition of m^6^A (19). Additionally, recently reported high-confidence m^6^A positions in *E. coli* (21), which were identified through a combination of DRS sequencing data analysis and the antibody-based MeRIP technique, could not be validated with SELECT in our hands or with DRS data analyzed using three different programs to assess the presence of m^6^A modifications in *E. coli* RNA (ELIGOS, Nanocompore, and CHEUI-diff m^6^A) (11,51,52). We compared DRS data from the WT with data from an IVT RNA control and the Δ*rlmF*Δ*rlmJ* mutant, the latter of which lacks the genes encoding m^6^A methyltransferases for the 23S rRNA. We detected the known m^6^A modification at position A2030 but not at A1618 in the 23S rRNA, consistent with a previous report (17). Notably, the m^6^A abundance in mRNA samples prepared using our protocol was ∼0.15 per 1000 As, which is less than 10% of the levels previously reported in *E. coli* (19). Nevertheless, our MS data support the idea that RlmF (an ortholog of METTL16) and RlmJ are at least partially responsible for m^6^A modifications in mRNA. This is plausible, given that bacteria lack a nucleus and thus all reactions occur in a single compartment, and was further supported by an *in vitro* methylation assay. Using a DRS approach, we were not able to detect any specific m^6^A sites in *E. coli* mRNA. This could be due to limitations in the sensitivity of the detection, the presence of m^6^A modifications in mRNAs with low abundance, low modification levels, population-level heterogeneity, and/or technical differences in growth conditions or sample preparations. It should be noted that the presence of m^6^A modifications in viral RNA genomes has recently been questioned (41). Previously, using immunoprecipitation and sequencing approaches, Chikungunya virus and dengue virus were shown to contain m^6^A modifications (72,73). However, a renewed investigation employing a combination of techniques such as SELECT, MS, and DRS could not confirm these earlier findings (41). Our work similarly emphasizes the need to use orthogonal methods to validate putative m^6^A positions identified with sequencing-based methods alone.

We analyzed the epitranscriptome of heat-stressed *E. coli* to test whether changes in RNA modifications could represent another level of regulation. At the same time, these experiments served as a validation of the use of DRS-based error rates to measure changes in RNA modification levels. Our ONT data demonstrated changes in rRNA modifications, specifically in m^6,6^A at sites 1518 and 1519 (as confirmed by MS data), as well as m^6^A, m^5^C, and m^4^Cm. All changes occurred at positions that are structurally closely linked and belong to the decoding center. This suggests that the heat stress response is accompanied by a remodeling of the decoding center through increased abundance of m^4^Cm, m^5^C, m^6^A, and m^6,6^A modifications. Consistent with these results, mutants for *rsmF* and *rsmA* (which encode the methyltransferases responsible for m^5^C1407, m^6,6^A1518, and m^6,6^A1519 modifications) demonstrate reduced growth under heat stress (33).

Our ONT dataset also allowed analyses of heat-stress-induced changes in tRNA modifications, comparable to previous studies assessing the roles of tRNA modifications in *E. coli* under heat stress (26). Although the DRS signals in tRNAs were noisier than those in rRNAs, it was possible to detect and evaluate modifications in many (∼70%) of the total number of unique *E. coli* tRNAs. In fact, our approach outperformed previous studies that used methods based on the interference of nt modifications using reverse transcriptases (26,60). Our DRS data were here highly correlated (ρ = 0.83) with the results of the gold-standard modification detection technique, MS, outperforming previous protocols based on dimethyl sulfate (DMS) or mutational profiling (MaP) (26). This is likely because some modifications (such as Ψ and m^5^U) that produce little or no RT impairment can be easily identified with DRS. The discrepancies identified here between the sequencing and MS results for Gm, m^6^A, and Cm may have occurred because varying levels of these modifications produced only small corresponding changes in ESB values (17). In contrast, modifications such as Ψ, m^6,6^A, m^2^G, Um, and m^1^G show large changes in ESB values in response to stoichiometric differences (17).

Although the averaged results from both ELIGOS and MS data did not indicate significant changes in D levels as a result of heat stress, decreased D was observed in specific tRNA positions at 45 °C (**Supplementary Figure 8**). This may be due to the role of D in enhancing tRNA structural flexibility (74); elevated temperatures inherently increase tRNA flexibility, and additional flexibility from D modifications at high temperatures could therefore be unfavorable. A phenomenon consistent with this hypothesis has been observed in *Bacillus*, in which the absence of D shows detrimental effects on bacterial growth at low temperatures (75).

Modification levels in the anticodon loop were here found to be generally reduced at 45 °C. For example, the DRS data revealed a significant decrease in Cm and Um levels at position 32 of the tRNA (**Figure 7C, Supplementary Figure 8**), which is consistent with previous reports of *E. coli* under heat stress (26). A significant decrease was also found in the level of the Ψ modification at positions 35, 38, 39, and 40 of the tRNA (**Figure 7C, Supplementary Figure 8**). These differences could not be assessed previously because mutational profiling (MaP) does not detect Ψ, and MS data lack the single-nt specificity enabled by DRS. Modifications in the anticodon loop are known to affect the decoding process through rare codon usages, translation frameshift, or wobble to maintain the reading frame (4,76). Decreased modification levels in the anticodon loop can result in translational infidelity, protein misfolding, and activation of the unfolded protein and heat shock responses; cells may also overexpress hypomodified tRNA to alleviate protein homeostasis (77,78), which could be advantageous under heat stress. In addition, we hypothesize that lower modification levels in the anticodon loop compensate for increased 16S rRNA modification levels in the decoding center.

Finally, we assessed the overall distribution of modifications throughout the *E. coli* transcriptome using a combination of thresholds designed to maximize the detection of modified sites while keeping the error rate below 15%. This revealed the presence of 387 putative modifications outside the tRNAs and rRNAs (i.e., in the mRNAs, ncRNAs, and intergenic regions) in *E. coli* grown at 37 °C. Many of the putative mRNA modifications observed at 37 °C were present in genes associated with general metabolism and RNA processing, suggesting that a primary function of mRNA modifications in *E. coli* may be related to the regulation of RNA itself. Transcripts containing modifications that increased in abundance at 45 °C compared to 37 °C were enriched in GO terms related to cell wall biosynthesis and transmembrane transport, indicating a role of the epitranscriptome in the regulation of the stress response. This is broadly consistent with the phenomenon of global changes reported in the epitranscriptome of *S. cerevisiae* in response to heat stress (79).

In summary, we here present the first systematic study of RNA modifications in *E. coli.* Deciphering a bacterial epitranscriptome requires a combination of complementary techniques, which here included DRS of high-quality RNA, MS, single-nucleotide PCR, and methyltransferase assays. These integrated approaches allowed us to simultaneously detect numerous modification types across RNA biotypes, including rRNAs, tRNAs, and mRNAs. On this basis, we found that the epitranscriptome of *E. coli* changes under heat stress, providing an additional level of regulation to fine-tune bacterial heat-stress responses.

## ACKNOWLEDGEMENTS

This work was financially supported by the Deutsche Forschungsgemeinschaft (DFG, German Research Foundation): Project numbers 325871075 (SFB 1309) to KJ (A07), PG (A08), SK (A01), and 464582101 (JU270/21-1) to KJ. SRB acknowledges financial support by the Alexander von Humboldt Foundation. LVC acknowledges the support of the Graduate School Life Science Munich (LSM). We thank Bibakhya Saikia for helpful discussion of the manuscript.

## Notes

### Competing Interest Statement

The authors have declared no competing interest.

### Summary of Updates

We have added a complete new set of direct RNA sequencing data for E. coli exposed to heat stress. We discuss the observed changes in known rRNA and tRNA modifications during stress, identify putative modification sites in the mRNA at two temperatures (37 and 45 degrees celsius) and detect changes in the putative mRNA modifications in response to heat stress. This is the first comprehensive report of changes in the epitranscriptome of E. coli under heat stress providing evidence for a further level of regulation to fine-tune the adaptation of bacteria to stress. In addition, we address the recent publication by Tan et al., 2024, (https://doi.org/10.1093/nar/gkae601), which was published after our study was originally submitted for review. The authors of this study report the existence of high-confidence m6A modification sites in the mRNA that contradict our findings. Specifically, we analyzed several positions reported as mRNA modification sites by Tan et al. using an orthogonal experimental method. We could not confirm the presence of m6A at the positions reported by Tan et al.. However, m6A modifications were detected at known sites in the 23S rRNA positive control, demonstrating the validity of our used experimental method. A discussion of these findings has been integrated into the revised manuscript.

