## Supplementary Material for "Direct RNA sequencing of the *Escherichia coli* epitranscriptome uncovers alterations under heat stress"

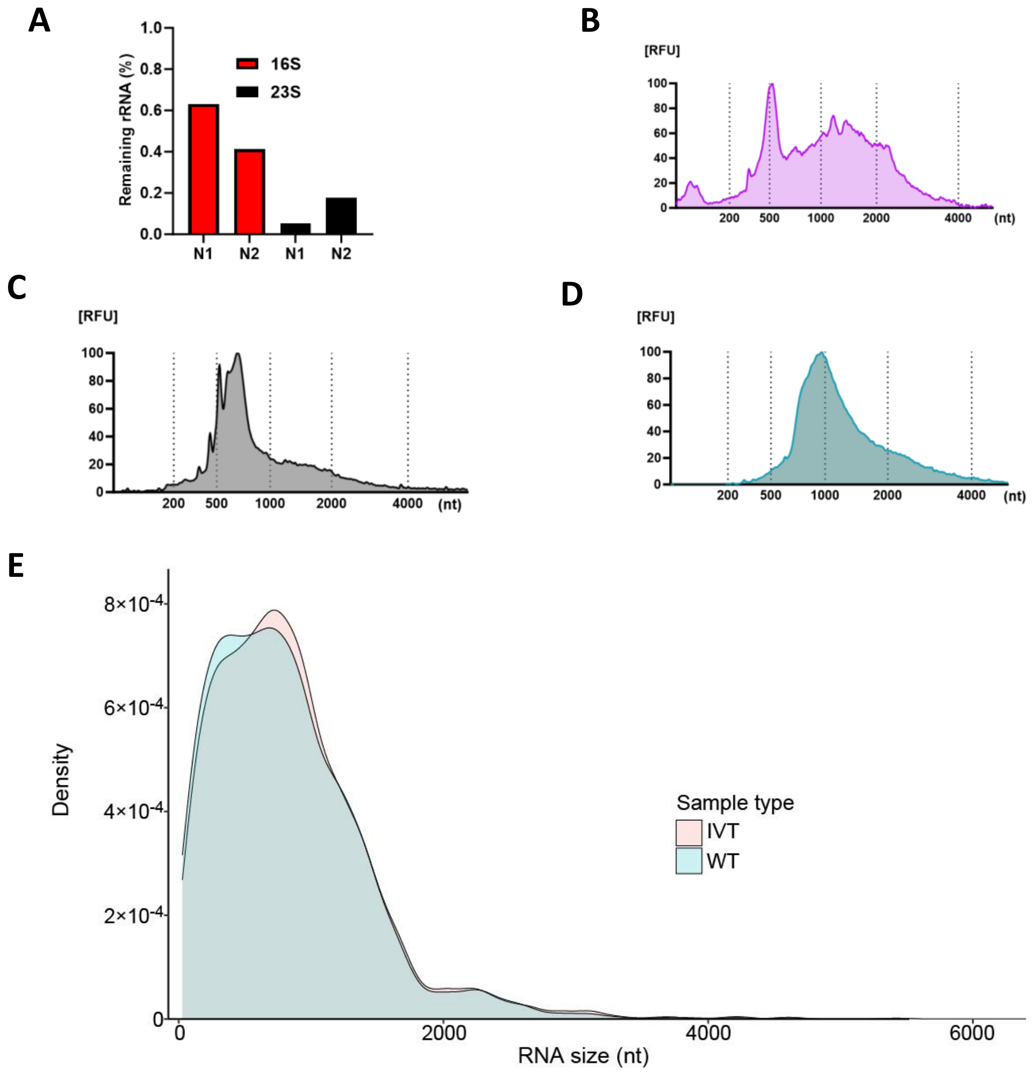


**Supplementary Figure 1. Quality control of prepared RNA samples**. (A) qRT-PCR was used to determine the abundance of 16S rRNA (red) and 23S rRNA (black) remaining after tRNA and rRNA depletion of WT RNA. rRNA abundance was normalized to that of the *recA* transcript. N1 and N2 indicate biological replicate samples. (B–D) Electropherograms showing relative fluorescent units (RFUs) at each nucleotide (nt) position in (B) tRNA/rRNA-depleted and polyadenylated WT samples, (C) *in vitro* transcribed (IVT) *E. coli* RNA samples before polyadenylation, and (D) IVT *E. coli* RNA after polyadenylation. (E) Size distribution of RNAs that were present at a depth of < 20 reads in the WT and IVT samples. There was no statistically significant difference in mean RNA size between the two sample types ($p > 0.05$; Student's $t$-test).

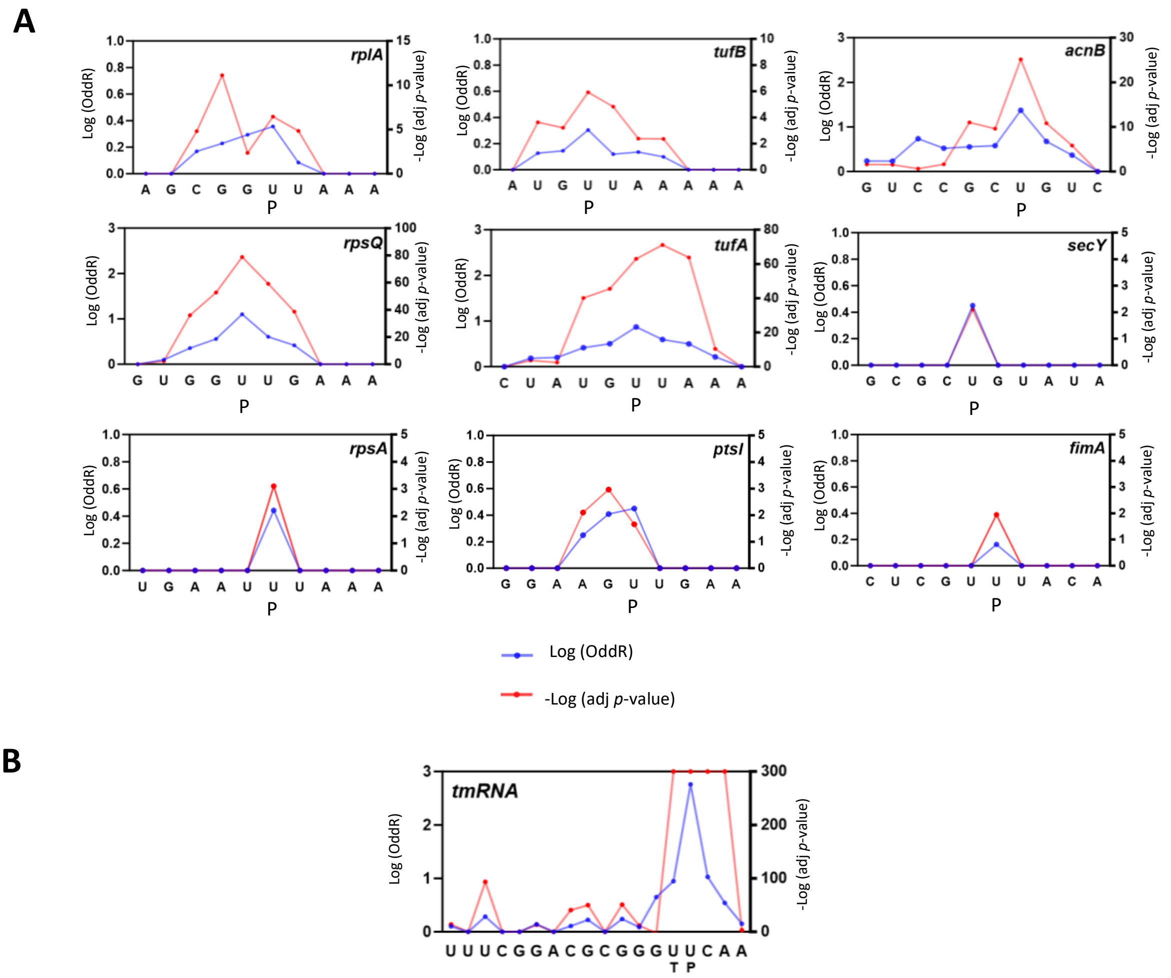


**Supplementary Figure 2. Detection of known RNA modifications in mRNA and transfer-messenger RNA (tmRNA).** (A, B) Log-transformed odds ratio (oddR) and negative log-transformed adjusted *p*-values for (A) nine sites previously identified as pseudouridine (P) in the *E. coli* mRNA [18] and (B) positions 341 and 342 of the *E. coli* tmRNA *ssrA*. T, 5-methyluridine ($m^5U$).

A

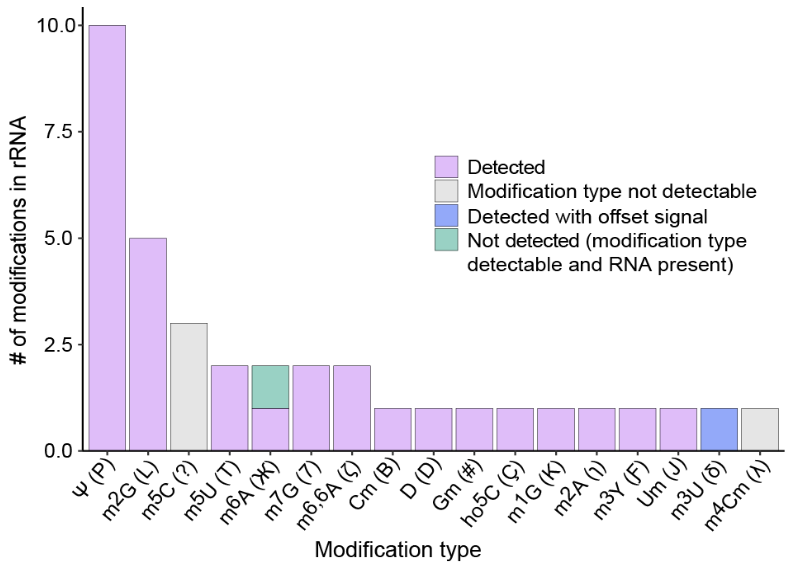


B

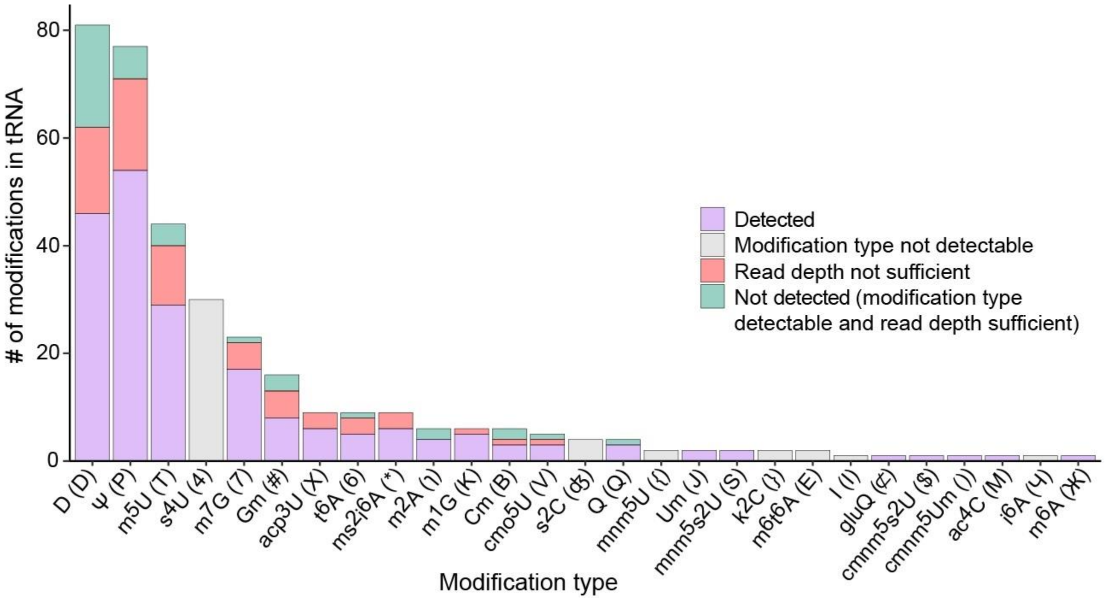


**Supplementary Figure 3. Detection of known modifications in *E. coli* tRNA and rRNA with ELIGOS.** The presence of known modifications in the ELIGOS output was evaluated for *E. coli* (A) rRNA and (B) tRNA. Modifications were classified as detected; of a modification type not known to be detectable with ELIGOS; contained in an RNA that was not present at a sufficient sequencing depth (20 reads); or of a modification type that was detectable with ELIGOS and contained in an RNA that was present at a sufficient sequencing depth in our dataset but not detected.

**A**

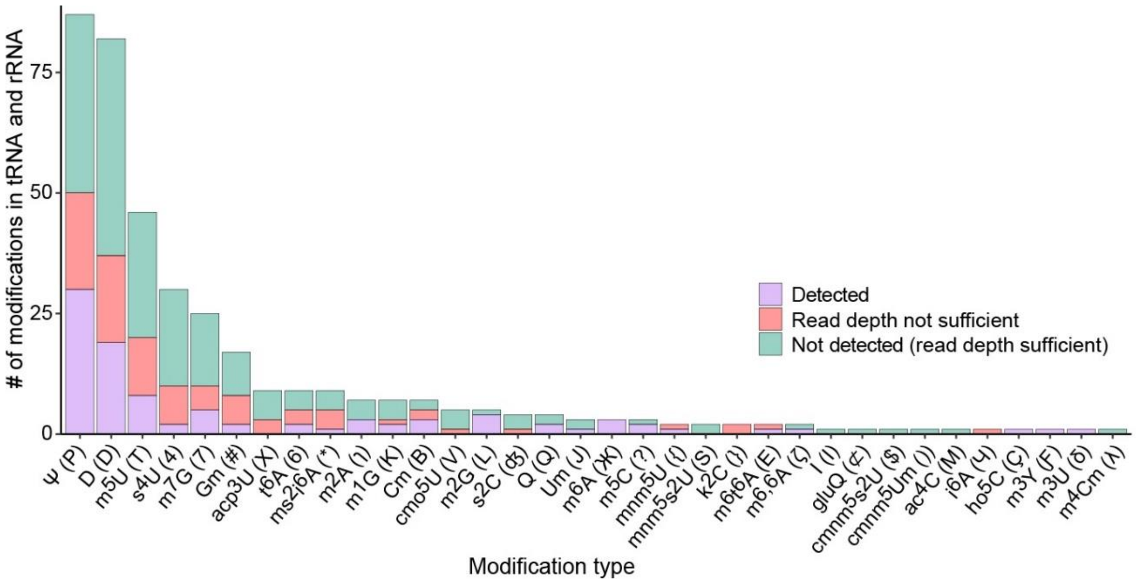


**B**

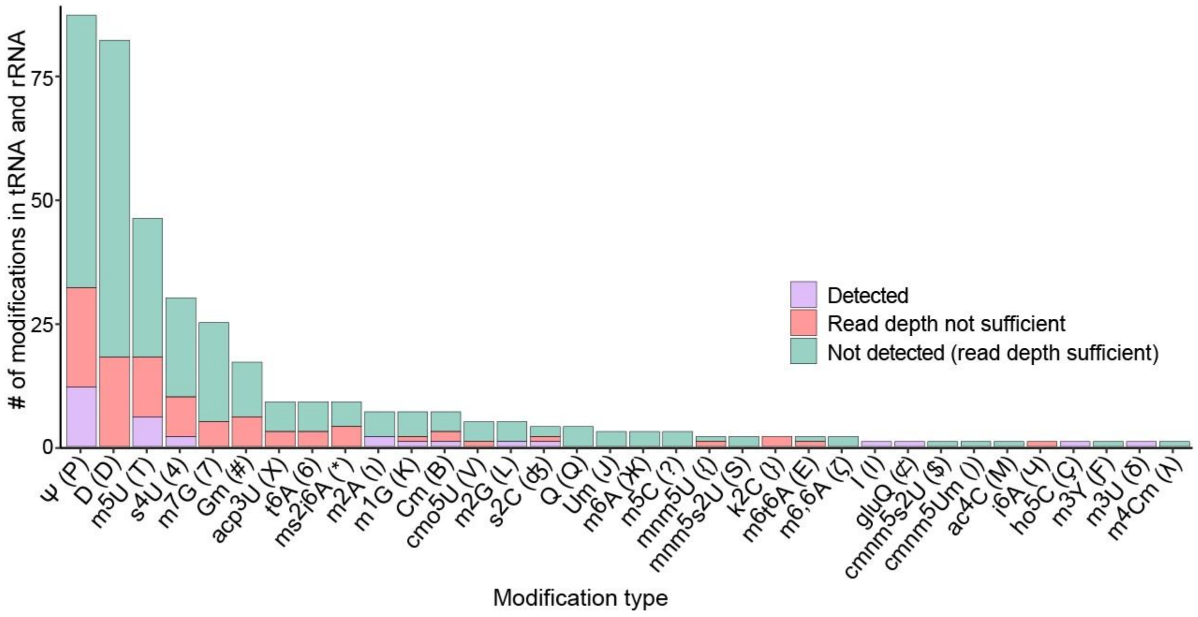


**Supplementary Figure 4. Detection of known modifications in *E. coli* tRNA and rRNA with Nanocompore and DRUMMER.** (A, B) Detection of known modifications in the *E. coli* tRNA and rRNA with (A) Nanocompore and (B) DRUMMER. Modifications were classified as detected; contained in an RNA that was not present at a sufficient sequencing depth (20 reads); or contained in an RNA that was present at a sufficient sequencing depth in our dataset but was not detected.

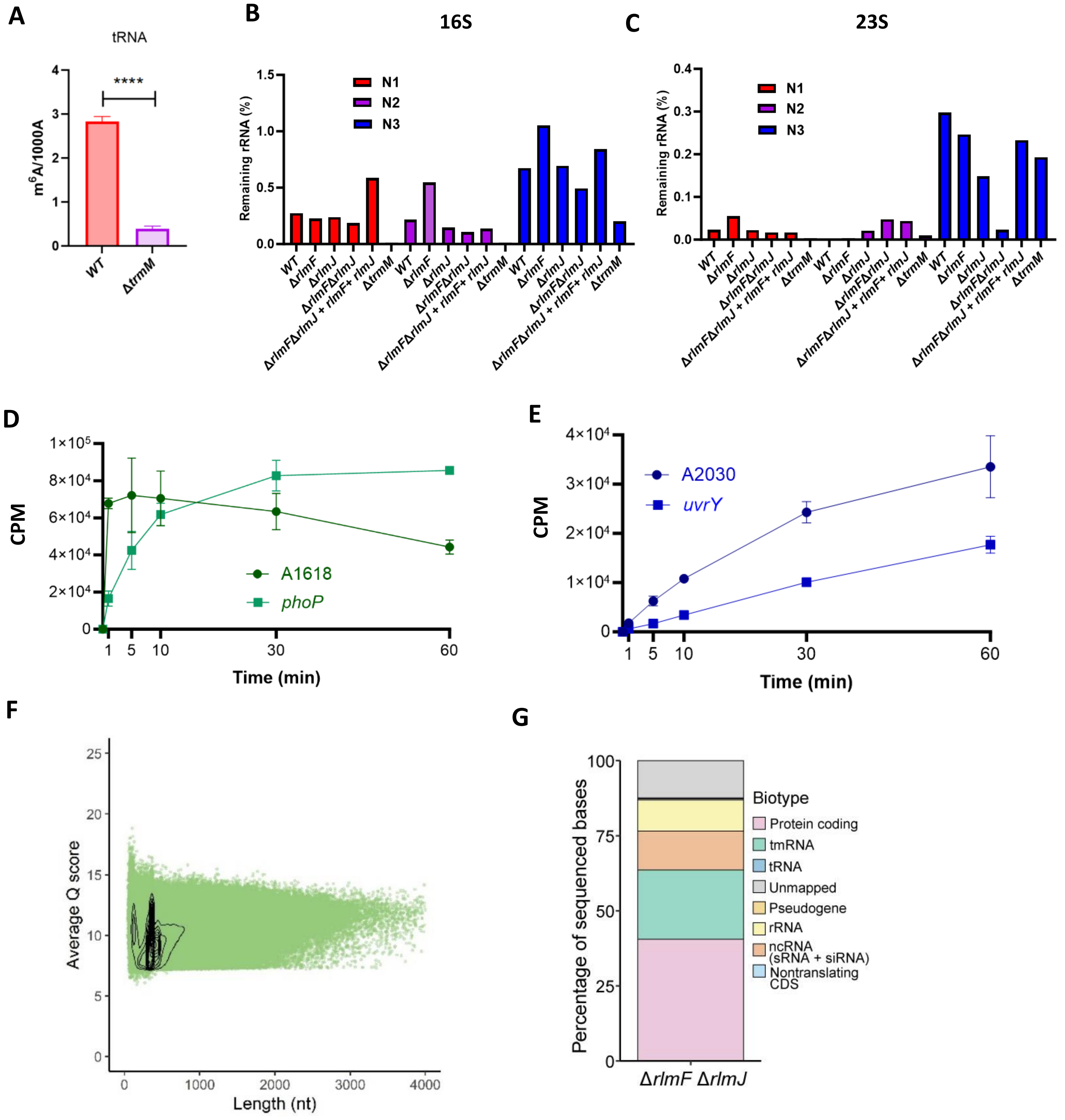


**Supplementary Figure 5. Detection of m$^6$A in WT and methyltransferase mutant *E. coli* RNA.** (A) Mass spectrometric measurements of tRNA samples collected from WT and Δ*trmM* mutant *E. coli* samples. Data are presented as the number of modified As (m$^6$A) per 1000 unmodified As (1000A). ****$p$ < 0.0001 (unpaired Student's *t*-test). (B, C) Percentages of (B) 16S and (C) 23S rRNA remaining after rRNA depletion in three biological replicates of several *E. coli* strains: WT; the single mutants Δ*rlmF*, Δ*rlmJ*, and Δ*trmM*; the double mutant Δ*rlmF*Δ*rlmJ*; and the complementation strain Δ*rlmF*Δ*rlmJ* + *rlmF* + *rlmJ*. qRT-PCR was performed to calculate the percentage of remaining rRNA (16S or 23S rRNA) in each sample, normalized to *recA* transcript abundance. N1–N3 indicate biological replicate samples. (D, E) Methyltransferase activities of (D) RlmF and (E) RlmJ over the course of 1 h. IVT RNAs of 100–300 nt in size were generated for *phoP* and *uvrY*, each of which contained an expected modification site; 30-nt RNAs containing As in the appropriate sequence context were also generated for the 23S rRNA positions 2030 and 1618. At time zero, the reaction was started by adding $^{3}$H-methionine. CPM, counts per minute. (F, G) Summary of sequencing data generated from Δ*rlmF*Δ*rlmJ E. coli* RNA, namely (F) average Q score per read vs. read length and (G) relative proportions of bases mapping to specific RNA types. tmRNA, transfer-messenger RNA; ncRNA, noncoding RNA; sRNA, small RNA; siRNA, small interfering RNA; CDS, coding sequence.

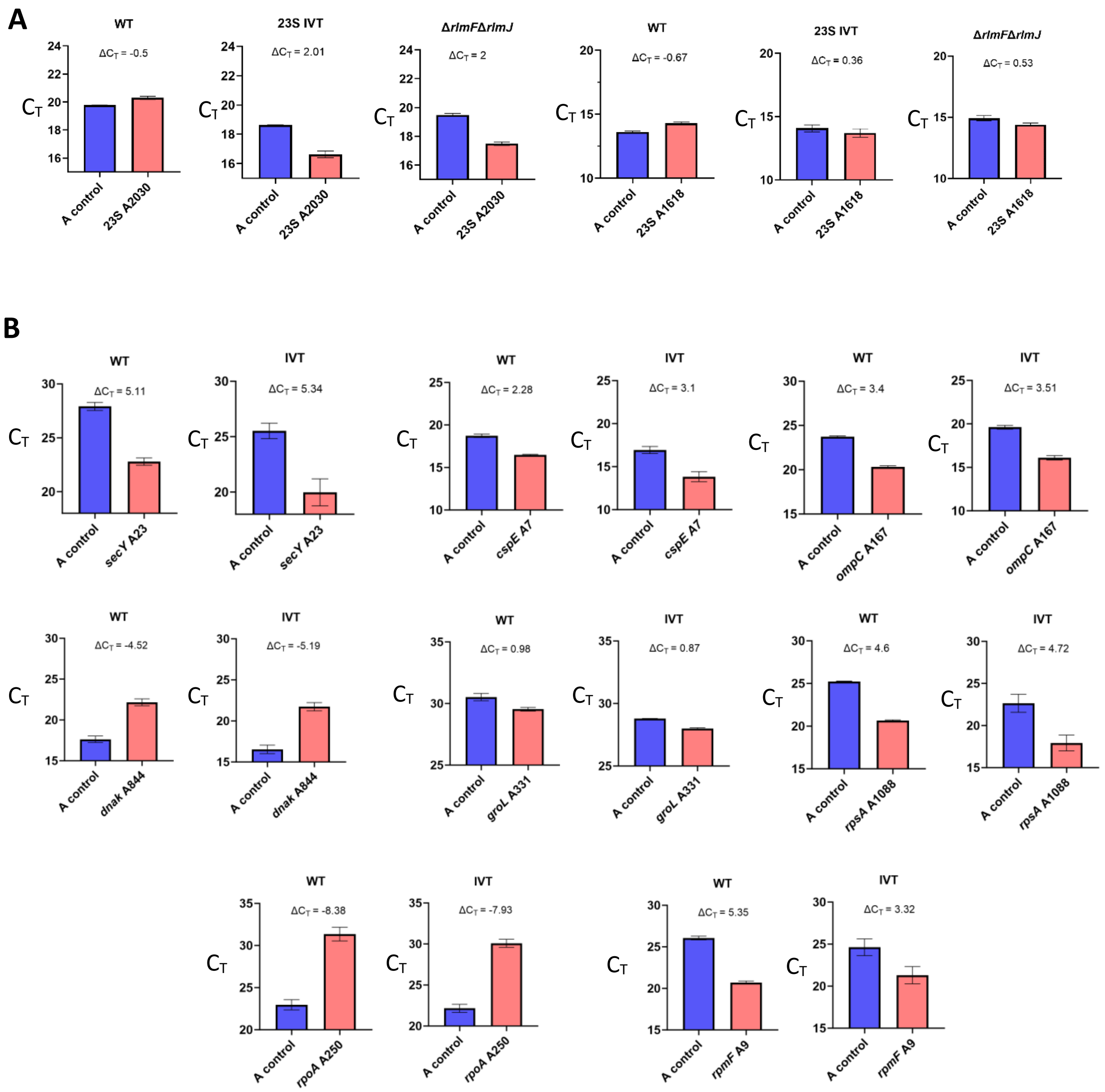


**Supplementary Figure 6. m$^6$A detection with SELECT**. (A) Detection of m$^6$A modifications at the A2030 and A1618 sites of the 23S rRNA in WT, IVT, and Δ*rlmF*Δ*rlmJ* *E. coli* with SELECT. (B) Assessment of eight *E. coli* mRNA sites previously identified as m$^6$A modifications [21] with SELECT. For each target m$^6$A site, a nearby unmodified control sequence was also tested (marked "A control"). Data are presented as the mean ± standard deviation from three biological replicates.

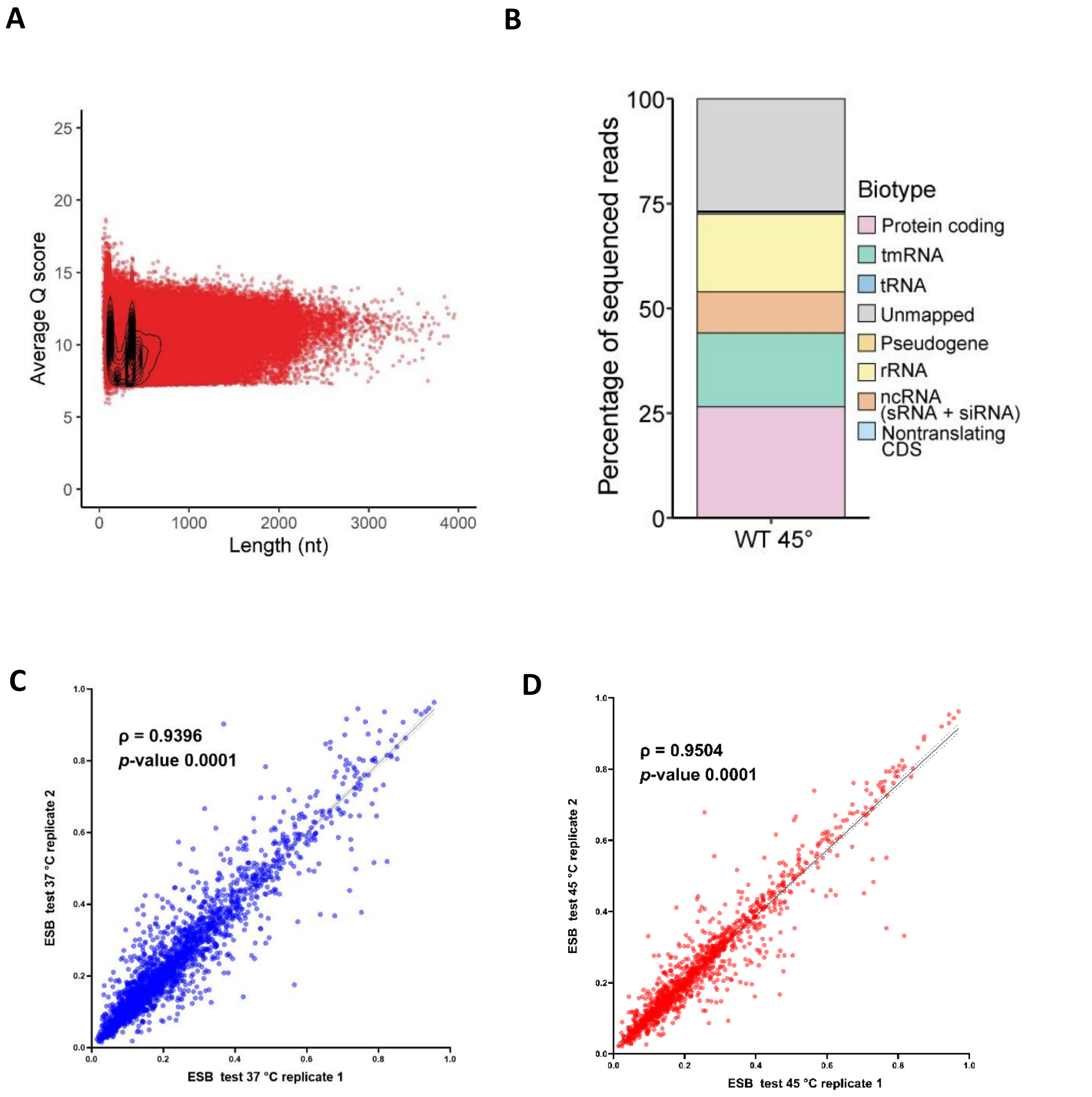


**Supplementary Figure 7. Summary of direct RNA sequencing data generated from *E. coli* exposed to heat stress.** (A) Average Q score per read vs. read length and (B) relative proportions of reads mapping to specific RNA types in samples extracted from WT *E. coli* grown at 45 °C. (C, D) Correlations between ESB test values from two biological replicates each (C) grown under standard growth conditions (37 °C) and (D) exposed to heat stress (45 °C). Correlations were calculated as Pearson's correlation coefficient (ρ).

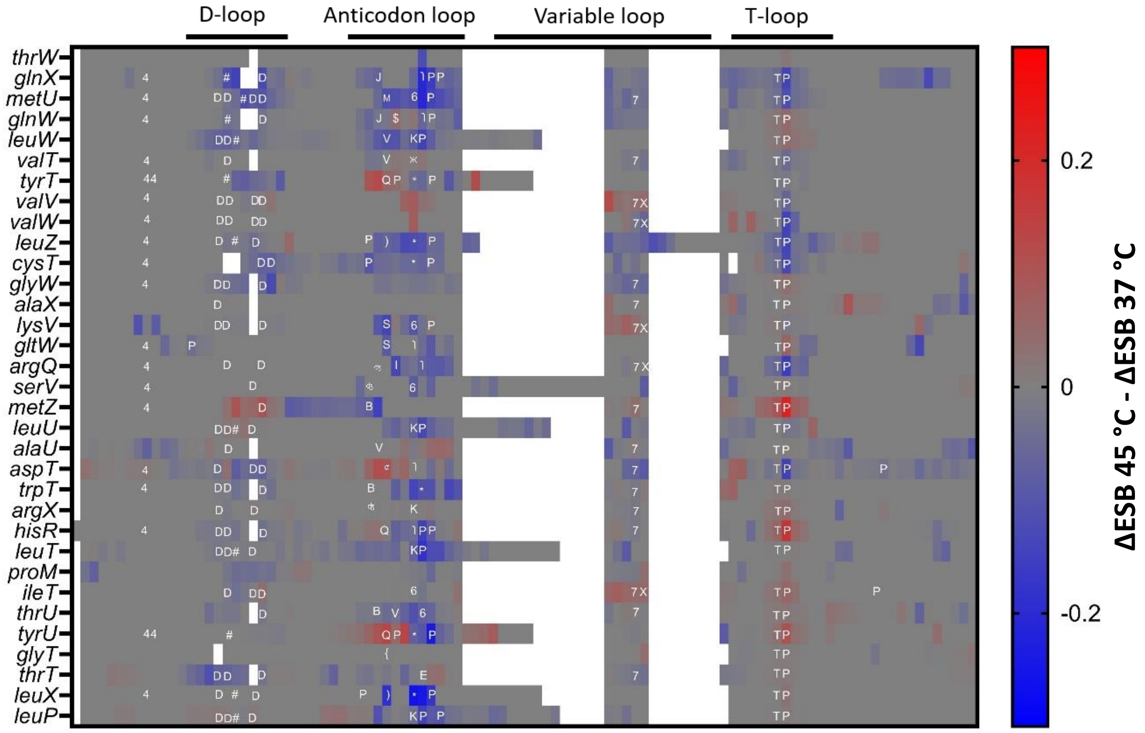


**Supplementary Figure 8. Changes in *E. coli* tRNA modification abundance in response to heat stress.** Changes in tRNA abundance were calculated as the ΔESB at 45 °C - ΔESB at 37 °C for positions across the 33 tRNAs containing modifications that were detected with ELIGOS.

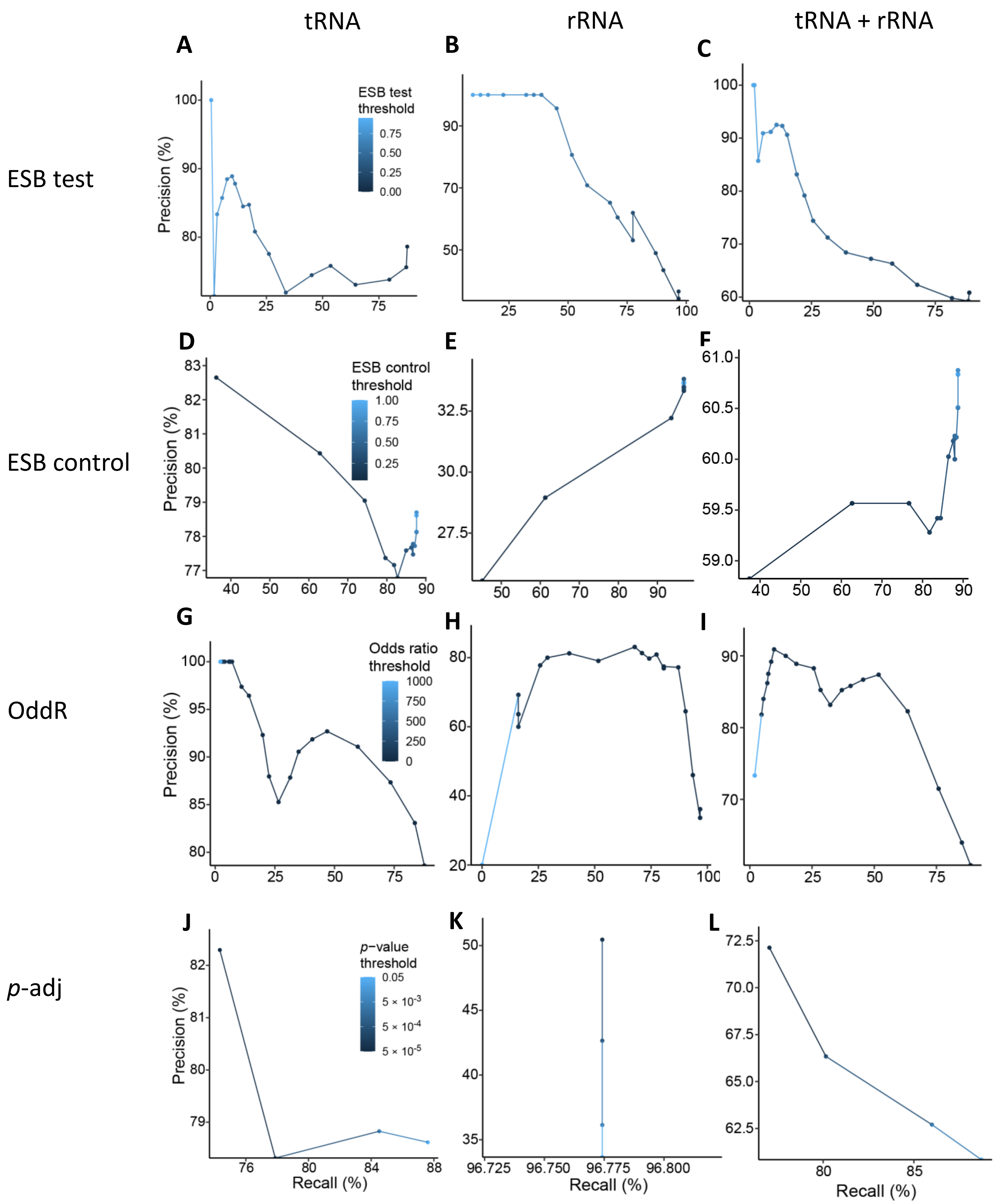

tRNA
rRNA
tRNA + rRNA
ESB test
ESB control
OddR
p-adj
A
B
C
D
E
F
G
H
I
J
K
L
ESB test threshold
ESB control threshold
Odds ratio threshold
p−value threshold
Precision (%)
Recall (%)

**Supplementary Figure 9. Precision and recall of known modification sites in *E. coli* tRNA and rRNA as detected with ELIGOS.** Modification sites in the *E. coli* rRNA and tRNA were detected with ELIGOS, then the results were examined using varying thresholds of the (A–C) percent error of specific bases (%ESB) in WT *E. coli* RNA (ESB test), (D–F) %ESB in the IVT control RNA (ESB control), (G–I) odds ratio (ESB test/ESB control) (oddR), and (J–L) adjusted *p*-value (*p*-adj). Left, tRNA only; center, rRNA only; right, tRNA + rRNA.

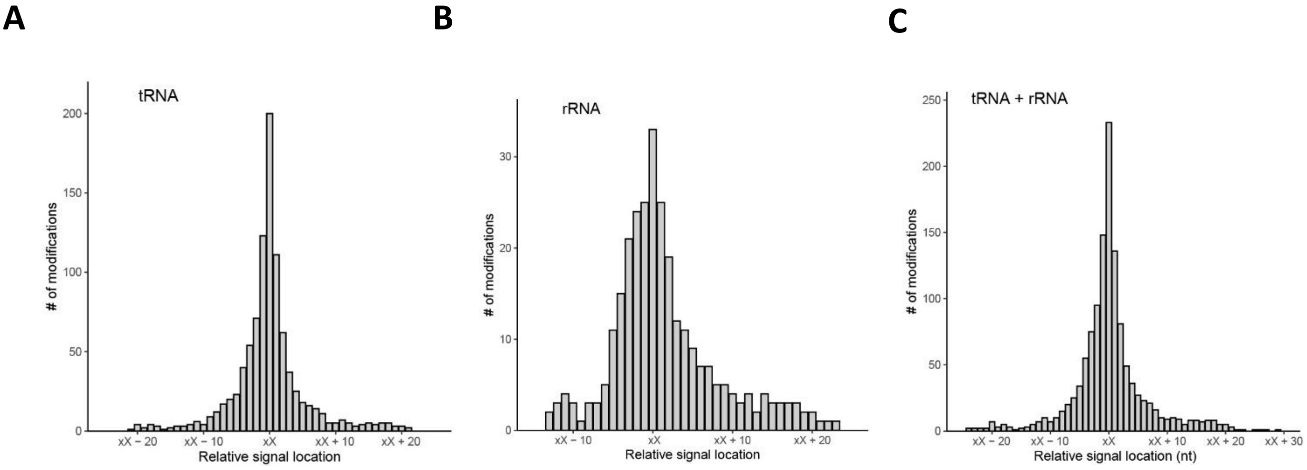


**Supplementary Figure 10. Distribution of modification signals around known modified sites**. The signals in the direct RNA sequencing data surrounding known modification sites (xX) were examined in the (A) tRNA only, (B) rRNA only, and (C) tRNA and rRNA combined.

**Supplementary Table 1.** Primers for mutant construction.

| Name | Sequence (5' to 3') |
|---|---|
| DEL_rlmF_for_f1 | GGCGCCAAGCTTCTCTGCAGGATCCGCCTGCGGAATTAATGAC |
| DEL_rlmF_rev_f1 | CTGGCGATTGACCGGTTTCTGGGCGGACATAGC |
| DEL_rlmF_for_f2 | CCGCCCAGAAACCGGTCAATCGCCAGCGTTAAAG |
| DEL_rlmF_rev_f2 | GCTAGCGAATTCGTGGATCCAGATGTGACTATCGGGCCGTTGAC |
| DEL_rlmF_chk_for | AGGCGAATGCAGTTACGTTG |
| DEL_rlmF_chk_rev | CGGCATGAACACTGGAGATTTG |
| DEL_rlmJ_for_f1 | GGCGCCAAGCTTCTCTGCAGGATCCTACCCAGGTGCTGTATTC |
| DEL_rlmJ_rev_f1_short | CCGCCAGTTCTTAGTGGCGATAACTGAGCATGG |
| DEL_rlmJ_for_f2_short | TTATCGCCACTAAGAACTGGCGGTACTGCCAGAC |
| DEL_rlmJ_rev_f2_short | GCTAGCGAATTCGTGGATCCAGATCATTTCTGGCCGTACATAGC |
| DEL_rlmJ_chk_for | GGTCGCGATACGCTTTATAC |
| DEL_rlmJ_chk_rev | TATTCCACGCCCGGAATATC |
| trmM Fw v2 | TGTATGGATATCGCTTCTCGGGTATTGTTCAATTAACCCTCACTAAAGGGCG |
| trmM Rv v2 | AAAGCCTGGGTCAGCGCCGTATACGCTTCGTAATACGACTCACTATAGGGCTC |
| c fw trmM | TTCGTCGAGTTCAAGTTCG |
| c Rv trmM | GATTATCCGGAACTGCTCAC |

**Supplementary Table 2.** Primers for reverse transcription quantitative PCR (RT-qPCR).

| Name | Sequence (5' to 3') |
|---|---|
| RT-qPCR recA Fw | CGGTTCGCTTTCACTGGATATCG |
| RT-qPCR recA rv | CCTGCAGCGTCAGCGTGGT |
| 23S Fw | TAACCGGCGATTTCCGAATG |
| 23S rv | CCAGACGCTTCCACTAACAC |
| 16S Fw | CAGGCGGTTTGTTAAGTCAG |
| 16S rv | CTTCGCCACCGGTATTCCTC |
| qPCRF for SELECT | ATGCAGCGACTCAGCCTCTG |
| qPCRR for SELECT | TAGCCAGTACCGTAGTGCGTG |

**Supplementary Table 3.** Mass transitions in dynamic multiple reaction monitoring mode with tandem quadrupole mass spectrometry (MS).

| Compound name | Precursor ion | MS1 Res. | Product ion | MS2 Res. | Ret time (min) | Delta Ret time | Fragmentor | Collision energy |
|---|---|---|---|---|---|---|---|---|
| **A** | 268,1 | Wide | 136 | Unit | 5,3 | 1 | 180 | 21 |
| **A SILIS** | 283 | Wide | 146 | Unit | 5,3 | 1 | 180 | 21 |
| **C** | 244,1 | Wide | 112 | Unit | 1,8 | 1 | 175 | 13 |
| **C SILIS** | 256 | Wide | 119 | Unit | 1,8 | 1 | 175 | 13 |
| **Cm** | 258,1 | Wide | 112 | Unit | 3,8 | 4 | 180 | 9 |
| **Cm SILIS** | 271 | Wide | 119 | Unit | 3,8 | 4 | 180 | 9 |
| **D** | 247,1 | Wide | 115 | Unit | 1,7 | 1 | 70 | 5 |
| **D SILIS** | 258 | Wide | 121 | Unit | 1,7 | 1 | 70 | 5 |
| **G** | 284,1 | Wide | 152 | Unit | 4,2 | 1 | 130 | 17 |
| **G SILIS** | 299 | Wide | 162 | Unit | 4,2 | 1 | 130 | 17 |
| **Gm** | 298,1 | Wide | 152 | Unit | 4,9 | 1 | 100 | 9 |
| **Gm SILIS** | 314 | Wide | 162 | Unit | 4,9 | 1 | 100 | 9 |
| **$m^1G$** | 298,1 | Wide | 166 | Unit | 4,9 | 1 | 105 | 13 |
| **$m^1G$ SILIS** | 314 | Wide | 177 | Unit | 4,9 | 1 | 105 | 13 |
| **$m^2G$** | 298,1 | Wide | 166 | Unit | 4,89 | 0,6 | 95 | 17 |
| **$m^2G$ SILIS** | 314 | Wide | 177 | Unit | 5,1 | 0,6 | 95 | 17 |
| **$m^3U$** | 259,1 | Wide | 127 | Unit | 4,7 | 2 | 75 | 9 |
| **$m^4Cm$** | 272 | Wide | 126 | Unit | 4,63 | 1 | 80 | 12 |
| **$m^5C$** | 258,1 | Wide | 126 | Unit | 3,6 | 1 | 185 | 13 |
| **$m^5C$ SILIS** | 271 | Wide | 134 | Unit | 3,6 | 1 | 185 | 13 |
| **$m^5U$** | 259,1 | Wide | 127 | Unit | 4,3 | 2 | 95 | 9 |
| **$m^5U$ SILIS** | 271 | Wide | 134 | Unit | 4,3 | 2 | 95 | 9 |
| **$m^{66}A$** | 296 | Wide | 164 | Unit | 7,2 | 1 | 130 | 21 |
| **$m^{66}A$ SILIS** | 313 | Wide | 176 | Unit | 7,2 | 1 | 130 | 21 |
| **$m^6A$** | 282,1 | Wide | 150 | Unit | 6,7 | 1 | 125 | 17 |
| **$m^6A$ SILIS** | 298 | Wide | 161 | Unit | 6,7 | 1 | 125 | 17 |
| **$m^7G$** | 298,1 | Wide | 166 | Unit | 3,5 | 2 | 100 | 13 |
| **$m^7G$ SILIS** | 314 | Wide | 177 | Unit | 3,5 | 2 | 100 | 13 |
| **$ms^2i^6A$** | 382 | Wide | 250 | Unit | 7 | 5 | 110 | 21 |
| **$ms^2i^6A$ SILIS** | 403 | Wide | 266 | Unit | 7 | 5 | 110 | 21 |
| **$t^6A$** | 413,1 | Wide | 281,1 | Unit | 6 | 1 | 130 | 9 |
| **$t^6A$ SILIS** | 434 | Wide | 297 | Unit | 6 | 1 | 130 | 9 |
| **U** | 245,1 | Unit | 113 | Unit | 3 | 1 | 95 | 5 |
| **U SILIS** | 256 | Unit | 119 | Unit | 3 | 1 | 95 | 5 |
| **Um** | 259,2 | Wide | 113 | Unit | 4,6 | 0,8 | 96 | 8 |
| **Um SILIS** | 271,1 | Wide | 119 | Unit | 4,6 | 0,8 | 96 | 8 |
| **Ψ** | 245,1 | Wide | 209 | Unit | 1,7 | 1 | 90 | 5 |
| **Ψ SILIS** | 256 | Wide | 220 | Unit | 1,7 | 1 | 90 | 5 |

Ret, retention; Res, resolution.

**Supplementary Table 4.** Primers for used for cloning.

| Name | Sequence (5' to 3') |
|---|---|
| RlmJ GA Fw | TTAACTTTAAGAAGGAGATATACCATGCTCAGTTATCGCCACAGC |
| RlmJ GA Rv | CTCGAGTGCGGCCGCAAGCTTCAACTCCGGCACGATCCAGCTTAC |
| pET 28 fw | TTGAAGCTTGCGGCCG |
| pET 28 rv | GGTATATCTCCTTCTTAAAGTTAAACAA |
| rlmF for NcoI | CATGCCATGGCAATGTCCGCCCAGAAACC |
| rlmF rev HIS HindIII | CGGGAAGCTTCTAGTGATGGTGGTGATGGTGTCCACCTCCACGCTGGCGA TTGACAAAACGGC |

**Supplementary Table 5.** Primers for *in vitro* transcription of selected genes.

| Name | Sequence (5' to 3') |
|---|---|
| A1618 | TCTACCTGACCACCTGTGTCGGTTTGGGGTCTATAGTGAGTCGTATTA |
| A2030 | GTACACTGCATCTTCACAGCGAGTTCCCTATAGTGAGTCGTATTA |
| T7-cas3 Fw | TAATACGACTCACTATAGGCTAGCTCATCCAGAACAA |
| cas3 rv | TACGCTGAACGATCGTC |
| T7-cysJ Fw | TAATACGACTCACTATAGTCTCTTCTGTTTTATGGGCG |
| cysJ rv | CAGCAGGCTGCTGATT |
| T7-rpoH Fw | TAATACGACTCACTATAGGATGTTGTCGGCTGAC |
| rpoH rv | GAACGGCGAAGGAGAC |
| T7-phoP Fw | TAATACGACTCACTATAGGCAGGACAAAGTCGAAGTATT |
| Phop rv | TTTGCCATTATTGCGTATC |
| T7-uvrY Fw | TAATACGACTCACTATAGACGTTCTACTTGTTGATGACC |
| uvrY rv | ACGGTAAGCATGATGATTTTG |
| T7-ppx Fw | TAATACGACTCACTATAGCACATGGTCATAGCCCG |
| ppx rv | ATACAGACGCTGGCAGG |
| T7-mazE Fw | TAATACGACTCACTATAGATGATCCACAGTAGCGTAAA |
| mazE rv | ACCAGTTCAGCAAGCGT |
| T7-dnaK Fw | TAATACGACTCACTATAGATCGCGGTTTATGACCT |
| dnak rv | GAGCGGAAGACAGTTCG |

**Supplementary Table 6.** Primers for single-base elongation- and ligation-based quantitative PCR amplification method (SELECT) to detect $N^6$-methyladenosine ($m^6A$) modifications.

| **Name** | **Sequence (5' to 3')** |
|---|---|
| 23S 2030 control down | 5PHOS/TCAATTTCACTGAGTCTCGcagaggctgagtcgctgcat |
| 23S 2030 control up | tagccagtaccgtagtgcgtgGCATCTTCACAGCGAG |
| 23S 2030 m6A down | 5PHOS/CACAGCGAGTTCAATTTcagaggctgagtcgctgcat |
| 23S 2030 m6A up | tagccagtaccgtagtgcgtgCGGGTACACTGCATCT |
| 23S 1618 control down | 5PHOS/GCTTAGAGGCTTTTCCTGcagaggctgagtcgctgcat |
| 23S 1618 control up | tagccagtaccgtagtgcgtgCGATTTGATGTTACCTGA |
| 23S 1618 m6A down | 5PHOS/GTGTCGGTTTGGGGTAcagaggctgagtcgctgcat |
| 23S 1618 m6A up | tagccagtaccgtagtgcgtgATTCTCTACCTGACCACC |
| cspE 7 m6a down | 5PHOS/AGACATCAAAATTACCTTTAcagaggctgagtcgctgcat |
| cspE 7 m6a up | tagccagtaccgtagtgcgtgCTTAACGTTACCTTTAATCT |
| cspE control down | 5PHOS/GAAACCGAATCCTTTGGACcagaggctgagtcgctgcat |
| cspE control up | tagccagtaccgtagtgcgtgTGCTGCCGTCTTCCGGAGTAA |
| dnak 844 m6A down | 5PHOS/AACGTCGGTCTGCTGAGCcagaggctgagtcgctgcat |
| dnak 844 m6A up | tagccagtaccgtagtgcgtgGCAGTGATGTATGGCAGGT |
| dnak control down | 5PHOS/GTGTTTCGGACCGGTCGcagaggctgagtcgctgcat |
| dnak control up | tagccagtaccgtagtgcgtgACGAGTCACTTTGATGTTCA |
| groL 331 m6A down | 5PHOS/GCCCGCAGCAACAGCTTTcagaggctgagtcgctgcat |
| groL 331 m6A up | tagccagtaccgtagtgcgtgTTTCAGGTCCATCGGGTTC |
| groL control down | 5PHOS/CGGGTTCATGCCCGCAGcagaggctgagtcgctgcat |
| groL control up | tagccagtaccgtagtgcgtgCGATACCACGTTTCAGGTCCA |
| ompC 167 m6A down | 5PHOS/AGGTCTGGTCGCCATCTACAcagaggctgagtcgctgcat |
| ompC 167 m6A up | tagccagtaccgtagtgcgtgCTTTGAAGCCAAGACGCATG |
| ompC control down | 5PHOS/GGTCAGTAACCTGAGTTTCAcagaggctgagtcgctgcat |
| ompC control up | tagccagtaccgtagtgcgtgGGCCGTAACCGGTCAGC |
| rpmF 9 m6A down | 5PHOS/ACGGCCATGGACCTTACTCcagaggctgagtcgctgcat |
| rpmF 9 m6A up | tagccagtaccgtagtgcgtgCGGGTTGGTTTATTCTGTTG |
| rpmF control down | 5PHOS/ACAGACAGCTGGTGACTCcagaggctgagtcgctgcat |
| rpmF control up | tagccagtaccgtagtgcgtgTTTTCACCAGAAGTTTTGTC |
| rpoA 250 m6A down | 5PHOS/GAGCAGGATTTCCAGGATATcagaggctgagtcgctgcat |
| rpoA 250 m6A up | tagccagtaccgtagtgcgtgCGCCAGCCCTTTCAGGT |
| rpoA control down | 5PHOS/AACTTCATCTTTGCCCTcagaggctgagtcgctgcat |
| rpoA control up | tagccagtaccgtagtgcgtgAGATTTATTCAAGGTAAGAA |
| rpsA 1088 m6A down | 5PHOS/TGTTGTGGGTTTCCGCGAACcagaggctgagtcgctgcat |
| rpsA 1088 m6A up | tagccagtaccgtagtgcgtgTACCTTCAACACGGTCGCCC |
| rpsA control down | 5PHOS/GTCAGACAGGTGAACCAGGcagaggctgagtcgctgcat |
| rpsA control up | tagccagtaccgtagtgcgtgCTGCAACGTTCCAGGAGA |
| secY control down | 5PHOS/AAGCCACCTTTGGCACTTTGAcagaggctgagtcgctgcat |
| secY control up | tagccagtaccgtagtgcgtgGTCTGCGTTTCAGCTCGCC |
| secY23 m6A down | 5PHOS/CTAATCCCGGTTGTTTAGCCcagaggctgagtcgctgcat |
| secY23 m6A up | tagccagtaccgtagtgcgtgCCACCTTTGGCACTTTGAAAA |

**Supplementary Table 7.** Primers for *in vitro* RNA transcription from total WT *E. coli* RNA.

| Name | primer sequence 5' to 3' |
|---|---|
| IVT Fwd Primer PCR | TAATACGACTCACTATAGCGAGGCGGTTTTCTGTTGGTGCTGATATTGCT |
| IVT Rev Primer PCR | ACTTGCCTGTCGCTCTATCTTC |
| 23S Fw T7 | TAATACGACTCACTATA GGTTAAGCGACTAAGCGTACA |
| 23S Rev T7 | AAGGTTAAGCCTCACGGTT |
| 16S Fw T7 | TAATACGACTCACTATAAATTGAAGAGTTTGATCATGG |
| 16S Rev T7 | TAAGGAGGTGATCCAACCG |
